# The novel nematicide chiricanine A suppresses *Bursaphelenchus xylophilus* pathogenicity in *Pinus massoniana* by inhibiting *Aspergillus* and its secondary metabolite, sterigmatocystin

**DOI:** 10.1101/2023.06.12.544558

**Authors:** Jiayu Jia, Long Chen, Wenjing Yu, Jun Su

## Abstract

**BACKGROUND:** Pine wilt disease (PWD) is responsible for extensive economic and ecological damage to *Pinus* spp. forests and plantations worldwide. PWD is caused by the pine wood nematode (PWN, *Bursaphelenchus xylophilus*) and transmitted into pine trees by a vector insect, the Japanese pine sawyer (JPS, *Monochamus alternatus*). Host infection by PWN will attract JPS to spawn, which leads to the co-existence of PWN and JPS within the host tree, an essential precondition for PWD outbreaks. Through the action of their metabolites, microbes can manipulate the co-existence of PWN and JPS, but our understanding on how key microorganisms engage in this process remains limited, which severely hinders the exploration and utilization of promising microbial resources in the prevention and control of PWD.

**RESULTS:** In this study we investigated how the PWN-associated fungus *Aspergillus* promotes the co-existence of PWN and JPS in the host trees (*Pinus massoniana*) via its secondary metabolite, sterigmatocystin (ST), by taking a multi-omics approach (phenomics, transcriptomics, microbiome, and metabolomics). We found that *Aspergillus* was able to promote PWN invasion and pathogenicity by increasing ST biosynthesis in the host plant, mainly by suppressing the accumulation of ROS (reactive oxygen species) in plant tissues that could counter PWN. Further, ST accumulation triggered the biosynthesis of VOC (volatile organic compounds) that attracts JPS and drives the coexistence of PWN and JPS in the host plant, thereby encouraging the local transmission of PWD. Meanwhile, we show that application of an *Aspergillus* inhibitor (chiricanine A treatment) results in the absence of *Aspergillus* and decreases the *in vivo* ST amount, thereby sharply restricting the PWN development in host. This further proved that *Aspergillus* is vital and sufficient for promoting PWD transmission.

**CONCLUSIONS:** Altogether, these results document, for the first time, how the function of *Aspergillus* and its metabolite ST is involved in the entire PWD transmission chain, in addition to providing a novel and long-term effective nematicide for better PWD control in the field.

## 1 INTRODUCTION

Pine wilt disease (PWD) has brought great economic and ecological damage to *Pinus* spp. forests and plantations worldwide ^1^. PWD is caused by the pine wood nematode (PWN) *Bursaphelenchus xylophilus*, which itself is transmitted by vector insects, namely the Japanese pine sawyer (JPS) *Monochamus alternatus* ^2–5^. Infection with PWN will turn the host into a lure that attracts JPS to spawn, this leading to the PWN and JPS co-existing within the host pine tree, an essential precondition for PWD transmission ^6^. In the last decade, tremendous effort has been expended towards developing effective strategies to impair the co-existence of PWN and JPS and thereby suppress the transmission PWD and potential outbreaks in stand. Microbes have proven substantial advantages for long-term pest control ^7–11^; however, their efficacy is strongly affected by local environments and a highly structured microbial community, as well as microbe-specific biochemical metabolites. Accordingly, the high-efficiency use of microbes varies with different conditions, being context-dependent ^12, 13^. Thus, exploring the biochemical basis enabling microbes to interfere with PWN and JPS co-existence is crucial for their use in effective long-term control of PWD.

Microbes are associated with PWN and play a crucial role in its invasion and pathogenicity ^1, 14^. The biological function of these PWN-associated microbes (PAMs) has been widely explored. Research has shown these PWN–PAM interactions can be beneficial (a mutualistic relationship), harmful (a parasitic/pathogenic relationship), or neutral ^1, 14–16^. These PAMs can also inactivate the PWN-induced resistance (mainly through the accumulation of reactive oxygen species [ROS]) in host pine trees, thus promoting the invasion and pathogenicity of PWN through adjustments to the microbial community or its members’ metabolites, including pathogenic factors, exoenzymes, and toxic secondary metabolites ^1, 6, 14, 17–20^. Yet, because of the complex structure and function of microbial communities, our understanding of the key microorganisms and their functions that affect the invasion and pathogenicity of PWN is still limited. This greatly restricts the exploration and utilization of potent microbial resources in the prevention and control of PWD.

After infection by PWN, the biosynthesis of volatile organic compounds (VOCs) is strongly augmented in host pine trees, which lures JPS for feeding and spawning ^21^. Several such VOC have been identified that are clearly capable of attracting JPS, including α-pinene, β-pinene, β-phellandrene, myrcene, and other terpene and sterols; some of them were successfully used to bait and trap JPS in the field ^22^. Previous studies have shown that PAM and host plant-associated microbes are directly or indirectly involved in the biosynthesis of the above VOCs via exoenzymes or metabolites ^23^. Hence, this hints that these PWN- and host-associated microbes might also affect the host preference of JPS, but this prospect warrants further exploration.

Based on above-mentioned lines of evidence, we hypothesized that a microbe can manipulate the co-existence of PWN and JPS through its metabolites, to influence PWD transmission. In our previous studies, by using metagenomic sequencing, we identified the key microorganism in host pine trees to be *Aspergillus*, whose abundance and activity are highly enriched during a PWN invasion and positively correlated with the amount and pathogenicity of PWN. Further, from correlations with metabolic data, it is evident that the pivotal metabolite of *Aspergillus* species, sterigmatocystin (ST), is highly induced by a PWN invasion, which might be the biochemical basis enabling *Aspergillus* to facilitate PWN invasion and pathogenicity. Given the above findings, the present study employed phenotypic, genetic, metabolic, and microbiological techniques to comprehensively clarify whether and how ST in the host plant *Pinus massoniana* enhances PWN invasion and pathogenicity; that is, by promoting host synthesis of VOC to attract JPS adults and ultimately sustaining the coexistence of PWN and JPS in their host, thus bolstering the transmission of PWD. Meanwhile, we speculated and tested whether the application of *Aspergillus* inhibitor (chiricanine A) could reduce the ST level in the host, to thus suppress PWN invasion and pathogenicity ^9^. The successful implementation of this project convincingly demonstrates the intrinsic biochemical mechanism of how key microorganisms, such as *Aspergillus* spp., are able to promote the transmission of PWD, which should encourage exploration of its application prospects in field settings.

## 2 MATERIALS AND METHODS

### 2.1 Plants and growth conditions

For the laboratory experiments, 2-year-old seedlings of *P. massoniana* were harvested from a nursery garden in a Shaxian Guanzhuang state-owned forest farm in Sanming, Fujian, China (26.5603°N, 117.7455°E). They were placed in growth chambers (light: dark = 16 h: 8 h, 70% humidity, 28 °C in light and 24 °C in dark) for 2 months before the experimental treatments began.

The field experiments were carried out in the same state-owned farm. This study area consisted of a 76-ha, 16-year-old pure *P. massoniana* plantation, where the annual average temperature and precipitation is 19.9 °C and 1375.2 mm, respectively.

### 2.2 Inoculation of pine wood nematode (PWN)

Adults of PWN were cultured as previously described ^9^. 1 mL of a PWN inoculation solution (5000 individuals/mL) was inoculated into *P. massoniana* seedlings as previously described, with 15 replicates (individual plants) used in each treatment. Double-distilled water served as the negative control.

### 2.3 Seedling injection with sterigmatocystin (ST), *A. arachidicola*, *A. sclerotioniger*, and chiricanine A

Sterigmatocystin (cat no. 10048-13-2, Shanghai Yuanye Bio-Technology Co., Ltd, Shanghai, China) was first dissolved in methanol, to prepare a stock solution (1mg/mL), then diluted to 1/10, 1/100, and 1/1000 with methanol as the working solution. These working solutions were separately injected into *P. massoniana* seedlings (1 mL per individual), as previously described ^24^. There was n = 15 per treatment, and a methanol solution served as the negative control. Plant samples were harvested at 14 days post-infection (dpi).

*Aspergillus arachidicola* (CBS117610) and *A. sclerotioniger* (CBS115572) were ordered from the Westerdijk Fungal Biodiversity Institute (https://wi.knaw.nl/). Each was inoculated separately into *P. massoniana* seedlings (1 mL per individual), with n = 15 for each treatment and double-distilled water used as the negative control.

Under laboratory conditions, 2-year-old *P. massoniana* seedling were inoculated with PWN, then injected with a solution of 4 ppm chiricanine A (cat no. B35217-5mg, Shanghai Yuanye Bio-Technology Co., Ltd, Shanghai, China) and emamectin benzoate (cat no. 155569-91-8, Merck, Shanghai, China). These solutions were injected into *P. massoniana* seedlings (1 mL per individual) as previously described ^24^; n = 10 for each treatment, for which a 4 ppm methanol solution served as the negative control. Plant samples were harvested at 14 dpi, then stored at –80 °C for subsequent experiments. In the field testing chiricanine A and 2% emamectin benzoate (EB) solution (PD20110688, Syngenta, Shanghai, China) were used for the trunk injections. The injection dose was based on the DBH of each tree (1 mL/cm). High-pressure trunk drilling and injection equipment (cat no. ZYJ15B, Greenman, Beijing, China) was used to apply the injections, by following a standard protocol (dose injected into stem at 1 m above the ground at a 45° angle).

### 2.4 Quantification of ROS accumulation in the host plant *P. massoniana*

The ST-infected and negative control *P. massoniana* seedling samples in the laboratory experiments were harvested at 14 dpi. Their collected pine needles were immediately placed in liquid nitrogen and stored in a –80 °C refrigerator. Then the amounts of ROS (cat no. ROS-1-Y, Comin Biology, Suzhou, China) and H_2_O_2_ (cat no. G0112W, Gris Biology, Suzhou, China) in these samples were measured by following the standard protocols of the above two kits.

### 2.5 Quantification of the relative PWN level

The amount of PWN in live plants was quantified by RT-qPCR (real-time quantitative polymerase chain reaction). Whole 2-year-old seedlings in the laboratory were used for the PWN quantification. The total genomic DNA of each plant sample was extracted using the MoBio PowerSoil DNA isolation kit (cat no.12855-50, MoBio, USA), according to the manufacturer’s protocol. The quantity and quality of DNA were measured on a NanoDrop 2000 photometer (Thermo Fisher Scientific, USA), with DNA integrity determined by 1% agarose gel electrophoresis. The extracted DNA was then stored at –80 °C until further use. Quantitative PCR were conducted using PWN-specific primers and a host-specific primer (one transcript from *P. massoniana* as the internal control) (Table S32), by using the Hieff^TM^ qPCR SYBR Green Master Mix (Low Rox Plus, cat no. 11202ES08, Yeasen, Shanghai, China) on a QuantStudio 6 Flex PCR (ABI). The qPCR signals were normalized to those of the reference gene *PST* in pine trees, by applying the 2^-ΔΔCT^ method ^25^. Biological triplicates with technical triplicates were used.

### 2.6. Quantification of plant defense genes and *aflR* of *Aspergillus* spp

Total RNAs were isolated from *P. massoniana* seedlings at 14 dpi by using the Trizol reagent (cat no. 15596026, Invitrogen, CA, USA). Each sample of total RNA (1 mg) was reverse transcribed by the PrimeScript^TM^ RT reagent Kit with gDNA Eraser (cat no. RR047A, Takara, Japan). The resistance genes examined were the same as those investigated in our previous study ^9^.

The *aflR* gene was extracted and searched by BLAST in the available genomes of the *Aspergillus* series at Genome/ NCBI (https://www.ncbi.nlm.nih.gov/genome). The sequences extracted from these genomes were aligned and consensus sequence used to design the primers online in Primer3 Plus (https://www.primer3plus.com). All the gene-specific primers used in this assay are listed in Table S32. RT-qPCR was carried out to quantify the gene expression level (as described in section 2.5).

### 2.7 Metabolome sequencing and analysis in the host plant *P. massoniana*

Sample preparation went as described in section 2.3, with three technical replications used. The metabolites were then extracted from each sample by following a previously described protocol ^26^. The Ultra High Performance Liquid Chromatography (UHPLC) separation was carried out using an A Dionex Ultimate 3000 RS UHPLC (Thermo Fisher Scientific, Waltham, MA, USA) equipped with an ACQUITY UPLC HSS T3 column (1.8 μm, 2.1×100 mm, 186009468, Waters, Milford, USA) by the Oebiotech Company (Shanghai, China). Set to a flow rate of 0.35 mL/min, the mobile phases were 0.1% formic acid in water (A) (A117-50, Thermo Fisher Scientific, Waltham, MA, USA) and 0.1% formic acid in acetonitrile (B) (A998-4, Thermo Fisher Scientific, Waltham, MA, USA). The column temperature was set to 45 °C, while the auto-sampler temperature was set to 4°C, and the injection volume was 5 μL ^27, 28^. Ensuing data were trimmed from different samples to distinguish the insect-induced metabolites. Next, commercial databases, including the Kyoto Encyclopedia of Genes and Genomes (KEGG; http://www.kegg.jp) and MetaboAnalyst (https://www.kegg.jp/) were utilized to search for ‘metabolitepathways’ (https://www.genome.jp/kegg/pathway.html).

### 2.8 Metagenome sequencing and analysis

For the microbiota within host pine tree, whole seedlings of *P. massoniana* (n = 5, containing PWN) after 14 dpi under the ST treatments were crushed in liquid nitrogen for their respective total DNA extraction, followed by metagenomic sequencing and analysis, as previously described ^9, 29^. Total genomic DNA was extracted from each sample by using the MoBio PowerSoil DNA Isolation Kit (12855-50, MoBio, United States) as per the manufacturer’s protocol. The DNA quantity and quality were measured on a NanoDrop 2000 spectrophotometer (Thermo Fisher Scientific, United States). This DNA was then sheared to 300-bp fragments by a Covaris ultrasonic crusher. To prepare each sequencing library, those fragments were treated by end repair, A tailing, and ligation of Illumina compatible adapters. Next all DNA sequencing libraries were deep-sequenced on an Illumina HiSeq platform at the Allwegene Company (Beijing, China). After every run, the image analysis, base calling, and error estimation were carried out using Illumina Analysis Pipeline v2.6. Quality control of the raw data, including the removal of adapter sequence and low-quality reads, was performed using Trimmomatic. High-quality sequences were compared with NR database and classified into different taxonomic groups, using the DIAMOND tool ^30^. Then MEGAHIT ^31^ was used to assemble the sequencing data, and the contigs were annotated with Prodigal software ^32^ to predict the open reading frames (ORFs). After that, CD-HIT software ^33^ constructed the non-redundant gene set. To compare the sequencing data with the non-redundant gene set, Bowtie ^34^ was used, after which the abundance information of genes in the different samples was counted.

### 2.9 Volatile organic compound (VOC) sequencing and analysis

For the VOC within the host pine tree, whole seedlings of *P. massoniana* (n = 5, with PWN) at 14 dpi under the ST treatments were examined, as previously described^25^. First, 500 mg (1 mL) of sample powder was transferred immediately into a 20-mL headspace vial (Agilent, Palo Alto, CA, USA) that contained an NaCl-saturated solution, to inhibit any enzyme reactions. These vials were sealed using crimp-top caps with TFE-silicone headspace septa (Agilent). During the Solid Phase Microextraction (SPME) analysis, each vial was placed accordingly at 60 °C for 5 min, then a 120 µm DVB/CWR/PDMS fibre (Agilent) was exposed to the headspace of a given sample for 15 min (also at 60 °C).

After completing that sampling procedure, desorption of VOCs from the fiber coating was performed in the injection port of the GC apparatus (Model 8890; Agilent), at 250 °C for 5 min, in the splitless mode. The identification and quantification of VOCs was carried out using an Agilent Model 8890 GC and a 7000D mass spectrometer (Agilent). The selected ion monitoring (SIM) mode was used for the identification and quantification of analytes by MS. The ensuing identified metabolites were annotated using KEGG Compound database (http://www.kegg.jp/kegg/compound/); annotated metabolites were then mapped to the KEGG Pathway database (http://www.kegg.jp/kegg/pathway.html). Pathways with significantly regulated metabolites mapped to them were then fed into MSEA (metabolite sets enrichment analysis), whose significance was determined by hypergeometric test’s *P*-values.

### 2.10 Insect host preference and diet quantification assay

Adults of JPS were collected from an experimental population reared at the Fujian Agriculture and Forestry University, Fuzhou, China. All experiments using them, as described below, were conducted in a growth chamber (light: dark = 12 h: 12 h, 70% humidity, 25 °C).

The host performance assay was done as described in a previous study^35^, albeit with some minor changes. Healthy and vigorous JPS were collected and starved for 24 h before starting the treatments, using a total 18 JPS adults per treatment (9 males, 9 females). Entire *P. massoniana* seedlings that had been inoculated with ST or 4 ppm methanol solution were the experimental odor source or negative control, respectively. Biological triplicates with technical triplicates were used. The performance of JPS on the host pine plants was expressed as an attraction ratio: [no. of choices by JPS / total no. Of JPS adults] * 100%. To determine statistical differences between treatment groups, first Chi-square goodness of fit test was applied, and then pairwise comparisons were made using multiple Mann-Whitney tests.

The diet of JPS adults was quantified as the consumed area of *P. massoniana* bark by these insects. Healthy and vigorous JPS individuals were collected and starved for 24 h before starting the treatments. A total of 12 JPS adults (6 males, 6 females) were used per treatment. Fresh 2-year-old *P. massoniana* seedlings from the various treatments were fed to the JPS for 3 days; then, the feeding area of the JPS was rubbed with transparent sulfuric acid paper, and this measured on grid coordinate paper. One-way analysis of variance (ANOVA; followed by Tukey’s test) was performed to determine the differences among the means of treatment groups.

All the JPS adults from the same treatment of each diet quantification assay were pooled into a single sample, crushed in liquid nitrogen, and divided into three equal aliquots for further analysis. The activities of exo-β-1,4-glucanase/cellobiose hydrolase (cat no. G0533W, Gris Biology, Suzhou, China), endo-β-1,4-glucanase (cat no. G0534W, Gris Biology, Suzhou, China), and β-glucosidase (cat no. G0535W, Gris Biology, Suzhou, China) in each sample were measured by following the standard protocols of corresponding kits. One-way analysis of variance (ANOVA; Tukey’s test) was implemented to determine the differences among the means of treatment groups.

## 3. RESULTS

### 3.1 Sterigmatocystin (ST) increases PWN pathogenicity by suppressing ROS accumulation in *P. massoniana*

Previous results shown that sterigmatocystin (ST) is highly positively associated with the pathogenicity of PWN. Here, we speculated that ST functions during the PWN invasion and pathogenic process (Fig. 1a). First, after PWN invasion, the ST level was highly induced (5.2 times, *P* < 0.01). Different concentrations (0.1 mg/mL, 0.01 mg/mL, 0.001 mg/mL) of the exogenous ST treatment applied to PWN-carrying *P. massoniana* (PCP) could significantly (*P* < 0.05) increase the PWN amount in the host, by 9.6, 6.6, and 4.6 times, respectively (Fig. 1b). Further, the death rate of host plants was measured at 2 months post-treatment with respect to different concentrations of ST (Fig. 1c). Evidently, the death ratio was positively correlated with the exogenous ST amounts; hence, ST could enhance the pathogenicity of PWN. We also measured the ROS level in PCP under the three ST treatments, finding that the rate of ROS (Fig. 1d) as well as H_2_O_2_ (Fig. 1e) production in PCP was significantly reduced by ST (*P* < 0.05) and negatively correlated (*P* < 0.01) with the exogenous ST concentration; this weakened resistance in PCP, thus benefiting the PWN invasion and pathogenic process. Further multi-omics (transcriptomics, microbiomes, and metabolic) data revealed that ST was capable of suppressing ROS accumulation through the regulation of a vast array of related genes (*c60547.graph_c0*, *c82953.graph_c0*, *c64867.graph_c0*, *c68789.graph_c0*, *c81022.graph_c0*), microbes (*Cladophialophora*, *Penicillium*, *Trichoderma*, *Achromobacter*, *Chitinophaga*, and *Flavobacterium*), and metabolites (maltotriose, 1-hexadecanoyl-2-(9Z-octadecenoyl)-sn-glycero-3-phosphoethanolamine and myo-inositol) (Fig. 1f). We also explored the ST-regulated metabolites from metabolic data. They were mostly enriched in arachidonic acid metabolism, flavonoid biosynthesis, phenylpropanoid biosynthesis, nucleotide metabolism, and *Cyprinus carpio* (common carp) pathways; this suggested secondary metabolites, ST inhibitors (stilbenoid), and VOCs (phenylpropanoids) were highly negatively (*P* = 0.021, *r* = −0.979), positively (*P* = 0.006, *r* = 0.994), and positively (*P* = 0.012, *r* = 0.989) correlated with the ST concentration, respectively (Fig. 1g). Also, the richness of several genera of microbes (*Enterobacter*, *Klebsiella*, *Pelagivirga*, *Herbaspirilum*, *Staphylococcus*, *Pseudovibrio*, *Rhizophagus*, *Cronobacter*, *Acinetobacter*, *Achromobacter*) was correlated with the ST concentration, and most of them were predicted to regulate the VOC and flavonoid biosynthesis (Fig. S1).

**Fig. 1.**
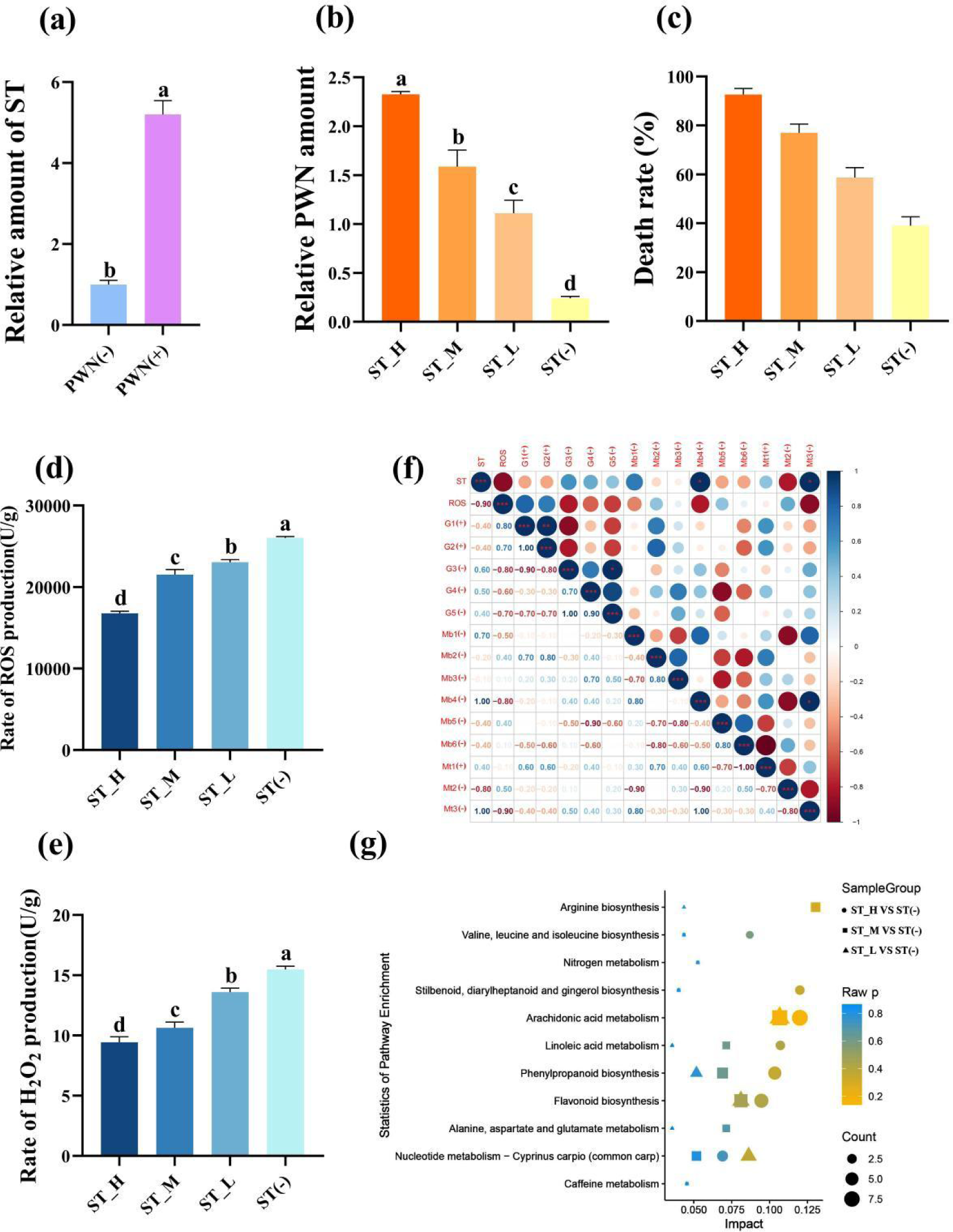
Sterigmatocystin (ST) increases Pinewood nematode (PWN) pathogenicity by suppressing ROS accumulation in *Pinus massoniana*. (a) Relative ST concentrations in the host plant *P. massoniana* quantified before (PWN(-)) and after (PWN(+)) invasion by PWN. The seedlings were inoculated with an equal amount of PWN (5000 individuals), followed by 0.1 mg/mL (ST_H), 0.01 mg/mL (ST_M), 0.001 mg/mL (ST_L) or a 0 mg/mL (ST(-)) of ST as the treatment, after which the (b) relative PWN amount, (c) death rate, (d) rate of ROS production, and (e) rate of H_2_O_2_ production were quantified at 14 days post-infection. Data shown is the mean ± standard deviation (SD). (f) Correlations between ST and their correlated ROS, transcripts, microbes, and metabolites. Metabolite pathways of *P. massoniana* as induced by ST with a highly correlated concentration. Different lowercase letters above bar columns show significant differences between treatments at *P* < 0.05, based on a one-way ANOVA, with multiple comparisons made using Tukey’s test. G1–G5 denote different transcripts (c60547.graph_c0, c82953.graph_c0, c64867.graph_c0, c68789.graph_c0, c81022.graph_c0); Mb1–Mb6 correspond to different microbial genera (*Cladophialophora*, *Penicillium*, *Trichoderma*, *Achromobacter*, *Chitinophaga*, and *Flavobacterium*); Mt1–Mt3 indicate different metabolites [maltotriose, 1-hexadecanoyl-2-(9Z-octadecenoyl)-sn-glycero-3-phosphoethanolamine, and myo-inositol]. (g) KEGG enrichment map of metabolic pathways of metabolites found significantly related to ST. The abscissa represents the impact of each pathway and the ordinate represents the pathways’ name. Impact is expressed as the ratio of the number of differential metabolites to the number of metabolites annotated in a given pathway. The circle represents ST_H vs. ST(-), the square represents ST_M vs. ST(-), and the triangle represents ST_L vs. ST(-), whose sizes indicate the number of differentially expressed metabolites contained in that metabolic pathway. Coloring represents the *P-*values for the enrichment analysis.

### 3.2 Sterigmatocystin (ST) increases VOC accumulation in *P. massoniana* to attract *Monochamus alternatus* (JPS)

VOCs have been proven to determine the host preference of JPS^36, 37^. Accordingly, here we first quantified the VOC amount of *P. massoniana* under each exogenous ST treatment. The latter significantly increased the total VOC amount within *P. massoniana* by 212, 182, and 105 times when compared to the negative control, respectively (*P < 0.001*; Fig. 2a). Also, for 10 VOCs (trans-anethole; acetophenone, 4’-hydroxy-; humulene; niacinamide; 4a(2H)-naphthalenol, octahydro-4,8a-dimethyl-,(4.alpha.,4a.alpha.,8a.beta.)-; 6-octen-1-ol,3,7-dimethyl-, (R)-; alpha-pinene; butanoic acid,3-hydroxy-3-methyl-; phenol; beta-myrcene), their amounts were positively correlated with the ST concentration, including three reported JPS-attracting VOCs (acetophenone, alpha-pinene, beta-myrcene) ^31, 38^ (Fig. 2b). These results led us to speculate whether the exogenous ST treatment can lure JPS adults. So we conducted an olfactory experiment, which showed that exogenous ST treatment can significantly (*P* < 0.05) promote the selective ratio of JPS adults to the host pine tree, and that ratio increased with a higher ST concentration (Fig. 2c). Host performance of insects were based on that this behavior is benefit to themselves ^39–41^. Furthermore, we also observed whether feeding on ST-treated *P. massoniana* could benefit the development of JPS The consumption area of JPS larvae feeding upon ST-treated *P. massoniana* increased significantly (*P* < 0.05) over time, and the body weight of the 3^rd^ instar along with the spawning ratio of JPS eggs were both significantly increased by 1.67 and 4.7 times vis-à-vis the negative control, respectively (Fig.2d–f). The EG (endo-1,4-β-D-glucanase), CBH (exo-β-1,4-D-glucanase) and β-GC (β-glucosidase) activity in the gut of the JPS adults that fed upon *P. massoniana* treated with ST were quantified as well, these increasing by 1.24, 1.21, and 1.54 times relative to the negative control, respectively (Fig. 2g–i). Altogether, these results suggested ST is able to increase the VOC accumulation in *P. massoniana*, thus attracting the JPS, whose feeding and development obviously improves when eating ST-treated *P. massoniana*, which can therefore manipulate the host preference of JPS.

**Fig. 2.**
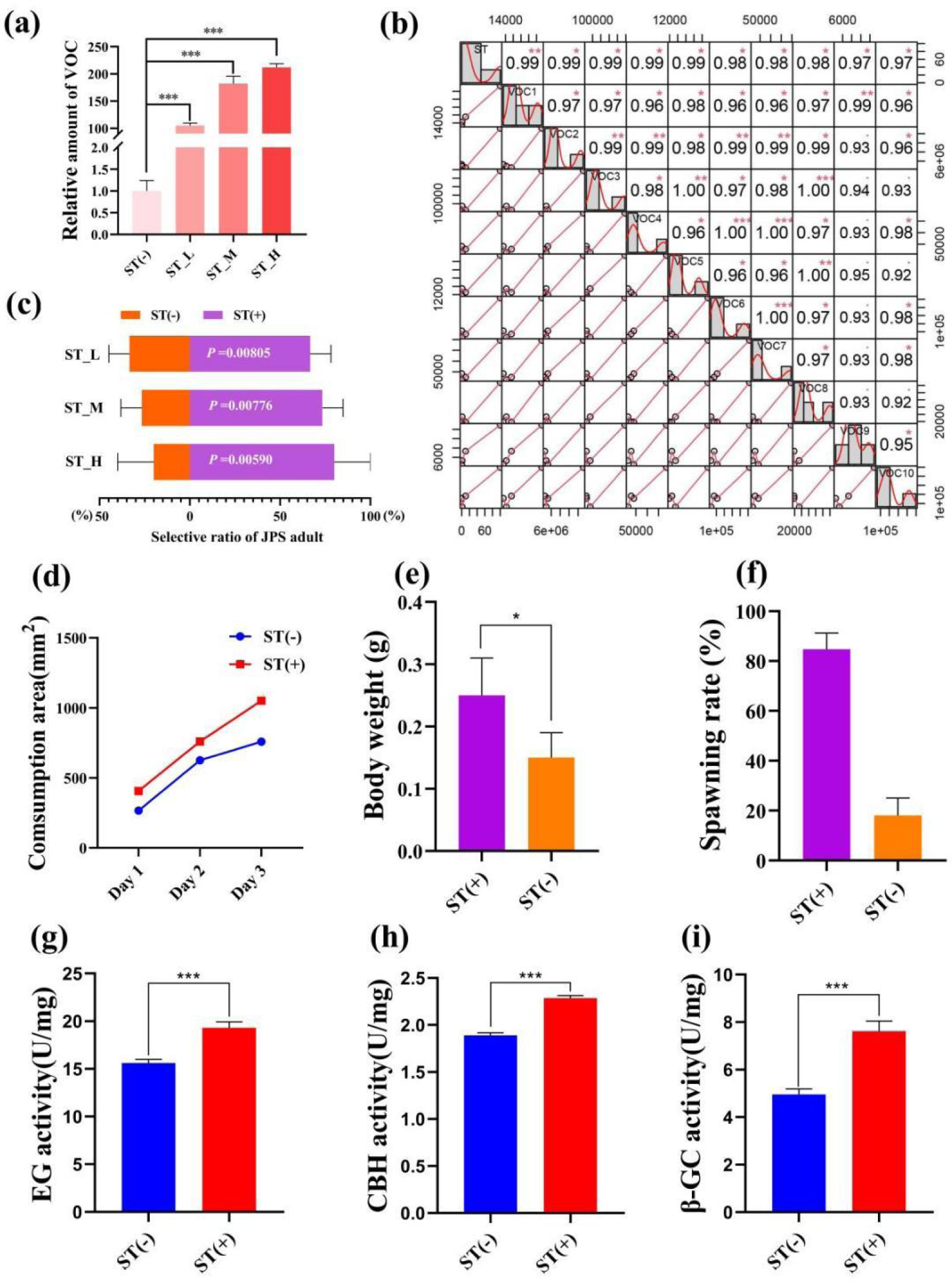
Sterigmatocystin (ST) increases VOC accumulation in *Pinus* Massoniana to attract *Monochamus alternatus* (JPS). (a) VOC amounts in host plant *P. massoniana* quantified 14 days after treatment with 0.1 mg/mL (ST_H), 0.01 mg/ ml (ST_M), 0.001 mg/mL (ST_L), or 0 mg/mL (ST(-)) of ST. (b) VOC compounds of *P. massoniana* induced by ST with a high concentration correlation. (c) Selective ratio of JPS to different plant samples were plotted, with the *P*-value for the *t* test between samples treated with (ST (+)) and without ST(ST(-)) presented in white. The (d) consumption area and (e) body weight of 3rd instar JPS larvae, and the (f) spawning rate, (g) EG (endo-1,4-β-D-glucanase) activity, (h) CBH (exo-β-1,4-D-glucanase) activity, and (i) β-GC (β-glucosidase) activity in the gut of the JPS adults that fed upon plant samples treated with (ST (+)) and without ST(ST(-)); data shown are the mean ± standard error (SE). The * and *** represents significant differences between treatments at *P* < 0.05 and *P* < 0.001, respectively, based on a one-way ANOVA, with multiple comparisons made using Tukey’s test. VOC1–VOC10 indicate different volatile organic compound (trans-anethole; acetophenone, 4’-hydroxy-; humulene; niacinamide; 4a(2H)-naphthalenol, octahydro-4,8a-dimethyl-,(4.alpha.,4a.alpha.,8a.beta.)-; 6-octen-1-ol,3,7-dimethyl-, (R)-; alpha-pinene; butanoic acid, 3-hydroxy-3-methyl-; phenol; beta-myrcene).

### 3.3 An associated microbe *A. arachidicola*, promotes PWN pathogenicity by increasing the sterigmatocystin (ST) accumulation in *P. massoniana*

Identify the vital effective microbe is crucial to achieving PWD control in the field. As an important metabolite in *Aspergillus*, ST is positively regulated by *alfR*. Although total species richness of *Aspergillus* harbored by the host plant was negligible influenced by PWN invasion and perhaps even slightly reduced (Fig. 3a), evidently the *alfR* expression level was significantly induced (14.06 times, *P* < 0.001) by PWN (Fig. 3b). We also found that only a few *Aspergillus* species (*A. arachidicola*, *A. fischeri*, *A. taichungensis*) were strongly (*P* < 0.001) increased or decreased (*A. sclerotioniger*, *A. awamori*, *A. aculeatus*) by PWN (Fig. S2). This prompted us to compare the functional differences between PWN-increased (*A. sarachidicola*) and -decreased (*A. sclerotioniger*) species and their associated microbes (Fig. 3c). After introducing it into *P. massoniana* via PWN, it was found that *A. arachidicola* significantly induced the accumulation of ST in the host (2.61 times, *P < 0.001*) whereas *A. sclerotioniger* did not (Fig. 3d). Further, the amount of PWN was respectively quantified in *A. arachidicola*- and *A. sclerotioniger*-infected PCP, which demonstrated the former can increase the PWN population size, whereas the latter cannot (Fig. 3e). Comparing the survival ratio of PCP infected by different microbes showed that it was sharply reduced after 2 weeks of infection by *A. arachidicola* but not *A. sclerotioniger* (Fig. 3f). Collectively, these results suggested that PWN-induced *Aspergillus* are sufficient to trigger ST accumulation that assists PWN invasion and pathogenicity.

**Fig. 3.**
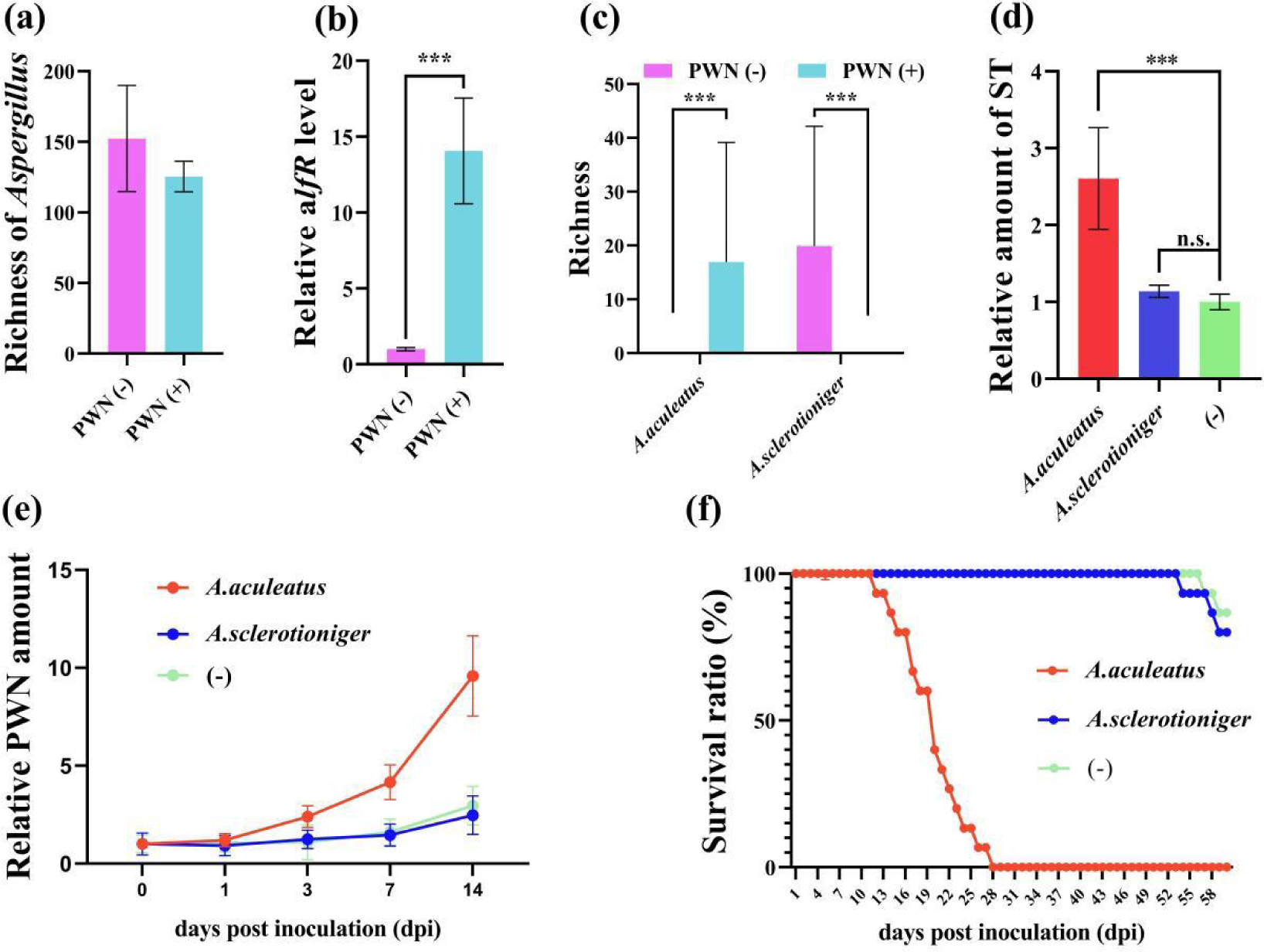
Associated microbe *Aspergillus arachidicola* promotes PWN pathogenicity by increasing the sterigmatocystin (ST) accumulation in *Pinus massoniana*. (a) Richness of microbes belonging to the *Aspergillus* genus, (b) expression level of ST synthesis promoting gene *aflR*, (c) richness of *A. arachidicola* and *A.* sclerotioniger in plant samples before (PWN(-)) and after (PWN(+)), invasion by PWN invasion are plotted. *Aspergillus arachidicola*, *A. sclerotioniger*, and inactivated *A. arachidicola* (-) were co-incubated with sterilized PWN, then inoculated into *P. massoniana* and the (d) ST amount, (e) relative PWN amount, and (f) survival ratio of different host plants were measured. Data shown are the mean ± standard deviation (SD). The *** represents significant differences between treatments at *P* < 0.001, based on a one-way ANOVA, with multiple comparisons made using Tukey’s test; ‘n.s.’ denotes no significant differences found.

### 3.4 The fungal inhibitor chiricanine A can suppress the species richness of *Aspergillus* fungi in *P. massoniana* host, thus curtailing in vivo PWN population size or pathogenicity in both laboratory and field tests

The *Aspergillus*-inhibitor chiricanine A has long been used to suppress ST accumulation in numerous crops^42^. Accordingly, we wondered whether chiricanine A could decrease the accumulation of ST by reducing the richness of PWN-induced *Aspergillu* species. We found that the application of chiricanine A significantly decreased (*P* < 0.001) the richness of PWN-induced *Aspergillu* spp. as well as the *aflR* expression level by 4 times and 1.77 times, respectively (Fig. 4a and b). When compared with emamectin benzoate, currently the most efficacious nematicide, chiricanine A was able to inhibit PWN for much longer (Fig. 4c). But when the host is inoculated with *A. arachidicola* or ST, either was clearly able to suppress that long-term effect of chiricanine A (Fig. 4d and e).

**Fig. 4.**
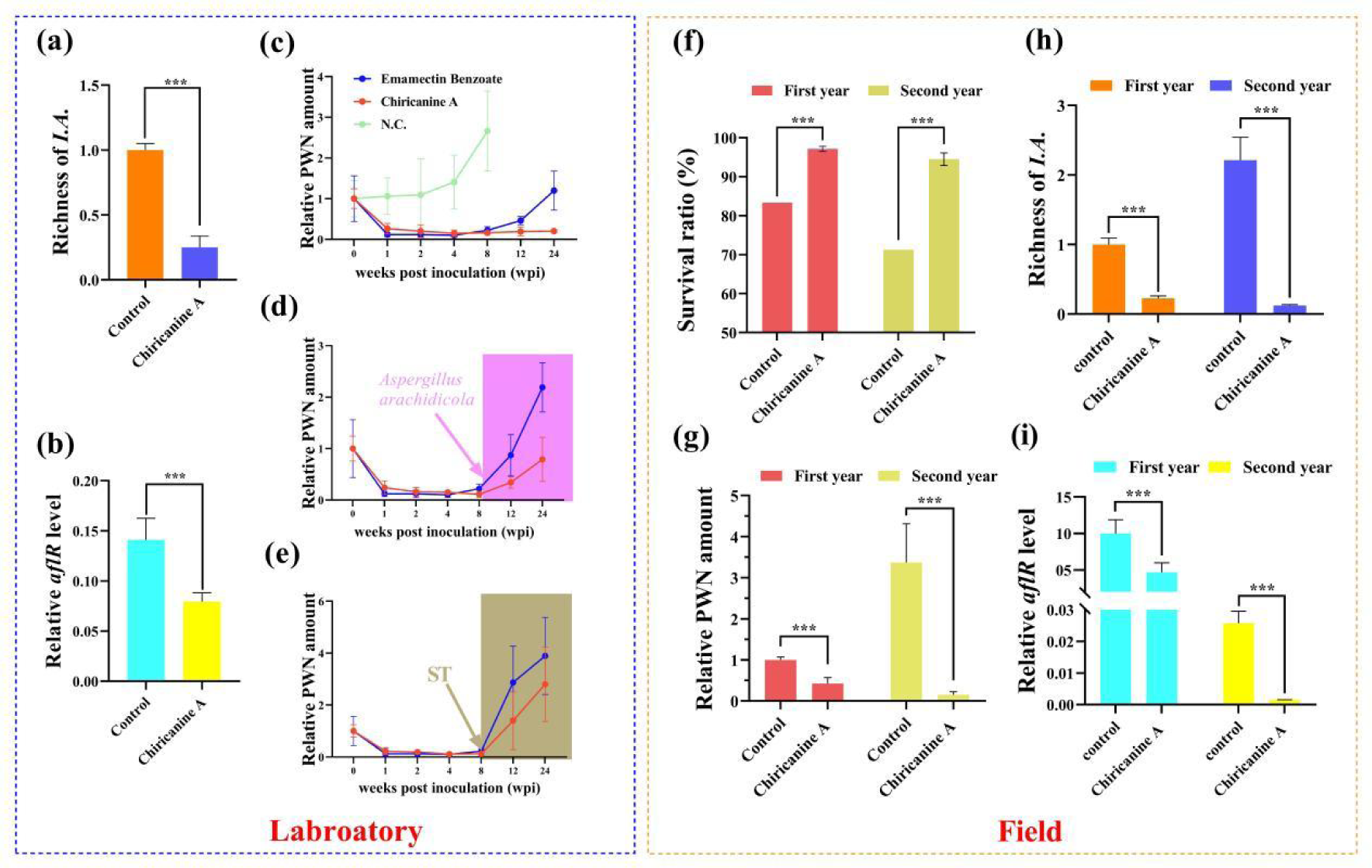
The fungal inhibitor chiricanine A can suppress the richness of *Aspergillus* fungi in *Pinus massoniana* to limit *in vivo* PWN population or pathogenicity, in both laboratory and field testing. Under laboratory conditions, 2-year-old *P. massoniana* seedlings were inoculated with PWN, and then inoculated with chiricanine A solution or methanol (control). For these samples, the (a) richness of *Aspergillus* fungi highly induced and (b) *aflR* expression level in response to PWN invasion were quantified. Relative PWN amount in PWN-carrying *P. massoniana* treated with (c) emamectin benzoate and chiricanine A, as well as those (d) injected with *A. arachidicola* or (e) ST at 8 weeks post-inoculation. In the field trial, a solution of chiricanine A or methanol (control) was injected into PWN-carrying *P. massoniana*, and its (f) survival ratio, (g) relative PWN amount, and the (h) richness of *Aspergillus* fungi highly induced and (i) *aflR* expression level in response to PWN invasion were quantified after the 1st and 2nd year post-injection. Data shown are the mean ± standard deviation (SD). The *** represents significant differences between treatments at *P* < 0.001, based on a one-way ANOVA, with multiple comparisons made using Tukey’s test.

We also tested the PWN-control functioning of chiricanine A in a 2-year field trial. The application of chiricanine A significantly increased the survival rate of PCP in both years vis-à-vis the control (1.17 and 1.33 times, *P* < 0.001) (Fig. 4f). Yet, significant reductions in the amount of PWN (2.36 and 22.05 times, *P* < 0.001), richness of PWN-induced *Aspergillus* spp. (4.44 and 18.07 times, *P* < 0.001), and the *aflR* expression level (2.15 and 18.07 times, *P* < 0.001) of PCP occurred in both years relative to the control (Fig. 4g–i). These results indicated that chiricanine A could serve as more efficient nematicide in the field by suppressing ST accumulation thereby performed a long-term control of PWD.

## 4. DISCUSSION

A paramount prerequisite to manipulating functional microorganisms is identifying those vital effective microbes that drive ecological phenomena, and elucidating their underlying biochemical mechanisms ^43^. Associated microbes enable linkages between PWN, host pine, and JPS ^14^; however, our understanding of how microbes directly affect the “PWN-host-JPS” complex through their produced metabolites are largely unknown ^1, 14, 44^. Here, we demonstrated that PWN-associated fungi, *Aspergillus* spp., are able to promote PWN invasion and pathogenicity by increasing biosynthesis of a secondary metabolite, sterigmatocystin (ST), in the host plant *P. massoniana*, mainly via suppressed ROS accumulation in hosts against PWN. Further, ST accumulation triggers VOC biosynthesis for attracting JPS to spawn and this drives the coexistence of PWN and JPS in host trees, thereby encouraging transmission of PWD (Fig. 5). Meanwhile, through the application of an *Aspergillus* inhibitor (Chiricanine A), we also showed that the absence of *Aspergillus* sharply restricts the development of PWN in *P. massoniana* (Fig. 4), further proving that *Aspergillus* is vital and sufficient to promote PWD transmission.

**Fig. 5.**
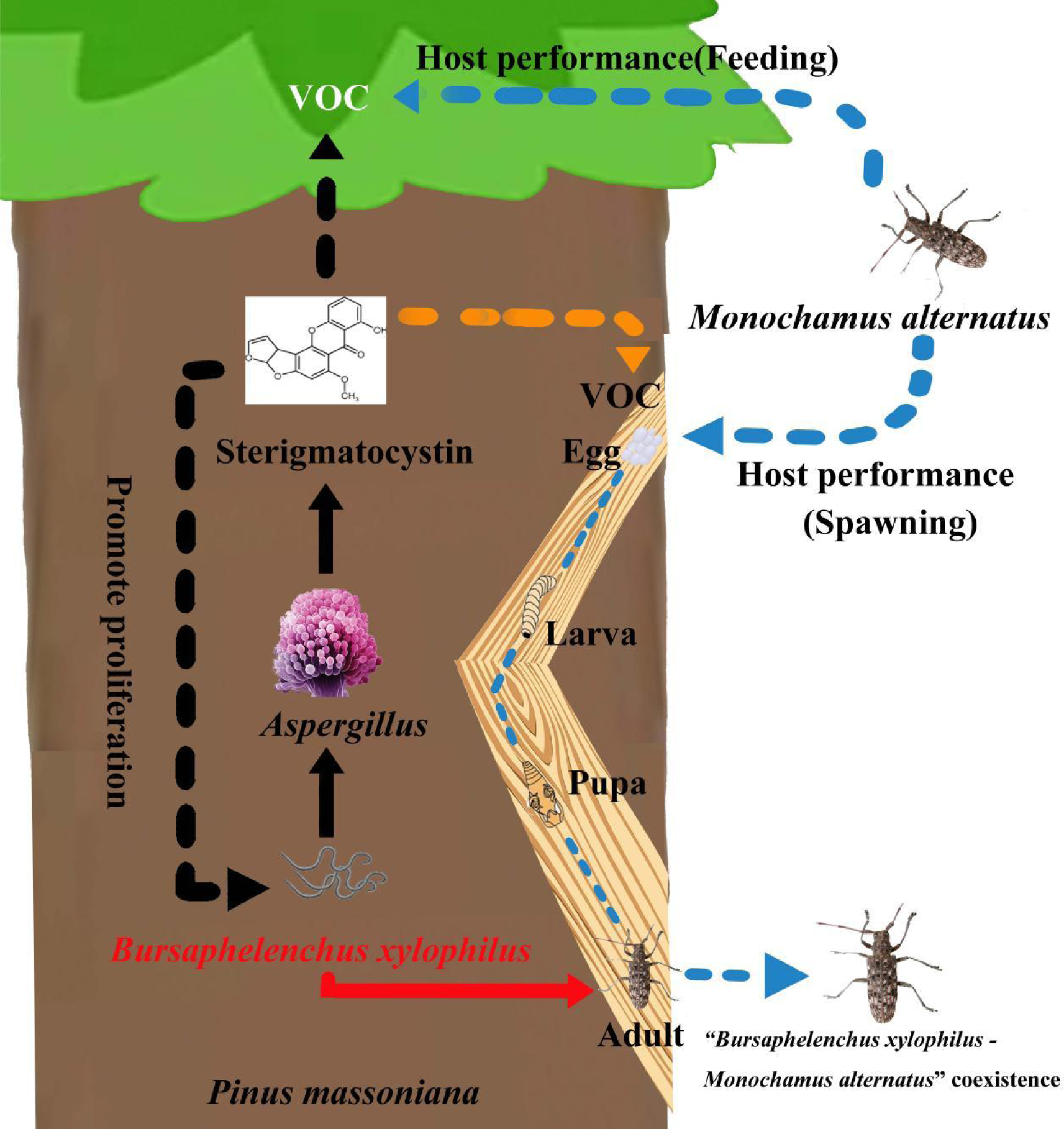
Schematic diagram of the pine wood nematode-*Monochamus alternatus* symbiosis promoted by *Aspergillus* fungi in *Pinus massoniana* host trees via sterigmatocystin (ST).

Ophiostomatoids and molds are two major fungal families well known for being highly correlated with the propagation and distribution of the PWN ^14^. Although molds do play crucial roles in multiple processes of PWD epidemics ^45^, in a way unlike the known contributions of ophiostomatoids to PWN (important food resources) ^6, 9, 46^, how molds are involved in PWD epidemics remains unclear. The present work, for the first time, clarifies how an important genus of molds, *Aspergillus*, contributes to the PWD epidemic via its secondary metabolite. The ST biosynthesis is conserved in *Aspergillus* ^47^ and promoted by *aflR*, and a pronounced positive correlation between ST and *aflR* expression was indeed observed in this study, but no correlation of ST with richness of *Aspergillus* spp. Although the absence of *Aspergillus* substantially restricts the PWN’s development in its host, among these identified *Aspergillus* species in the PWN-carrying *P. massoniana*, only three of them—*A. fischeri*, *A. arachidicola*, *A. taichungensis*—increased markedly in response to PWN invasion and were very likely transferred by PWN into host plants, in contrast to most of them (like *A. sclerotioniger*) decreasing considerably. These lines of evidence suggest PWN-associated *Aspergillus* fungi are able to reconstruct the host plant-associated microbial community to promote PWN development. This point is also supported by the diversity of the host plant-associated microbial community being severely reduced by the application of a low ST concentration yet increasing with higher ST concentrations. The above point can also explained by the various functions of *Aspergillus* species; most of the plant-associated *Aspergillus* are mutualists ^48–50^, one of the PWN-associated *Aspergillus* fungi, *A. arachidicola*, has been reported as being harmful to plant and beneficial for insect feeding ^48, 49^.

Despite ST being detectable in various of crops and foods ^51^, its functioning in plants has yet to be resolved. In this work, we demonstrated that ST was able to suppress ROS accumulation in the host pine tree via genetic, metabolic, and microbial level regulation processes. The richness of several genera of microbes—*Enterobacter*, *Klebsiella*, *Pelagivirga*, *Herbaspirilum*, *Staphylococcus*, *Pseudovibrio*, *Rhizophagus*, *Cronobacter*, *Acinetobacter*, and *Achromobacter*—was correlated with the ST concentration, and most of them were predicted to regulate the biosynthesis of VOCs and flavonoids. Further, these functional predictions were supported by the metabolic data. By quantifying the VOC amounts in host pine trees, we find that the total VOC amount is highly induced by ST (more than 100 times), yet only a few of them (mostly terpenes) are induced in a ST-dependent manner; this finding hints that ST might be a potential signal acting to regulate the host performance of insects. Although, some reported JPS-attracting VOCs (acetophenone, pinene, and myrcene) were among the above ST-induced metabolites, it was not clear whether these are all JPS-specific VOC, an aspect worth exploring further. Meanwhile, the amount of flavonoids is reduced substantially by ST, together with ROS accumulation being suppressed in host pine tree, implies that ST could also halt or paralyze the later stages of plant defense response ^52^. Plant flavonoids are widely proven to be toxic to pests, including PWN ^53, 54^. This suggests that PWN may have a highly effective detoxification system to withstand flavonoids and survive in the host plant by hijacking microbes. Further investigation of the PWN detoxification system acting against plant secondary metabolites, offers an excellent opportunity to explain how PWD outbreaks arise. Interestingly, we also found that synthesis of an ST inhibitor, stilbenoid ^47^, was promoted by ST, suggesting that plants may have a conserved system to respond to ST as a signal. Together, we hypothesize that ST can suppress the early stage of PWN resistance and reconstructs the microbial community in the host plant; it may also strongly activate the later stage of pest resistance (flavonoid accumulation) to limits competitors in the same niche, thereby promoting outbreaks of PWD in pine stands.

Emamectin benzoate (EB) is widely regarded as the most useful nematicide against the PWN, largely because it is environmentally friendly and has longer field residence time (i.e., at least 3 years) ^55, 56^. However, continuous injection is required to maintain an effective dose of EB in a pine tree or whole stand ^56–59^. Although neither EB nor chiricanine A can completely eradicate the PWN population in host pines, we propose that chiricanine A has a PWN control efficiency on par with EB, but is able to suppress the PWN population for a longer time. As a plant metabolite, chiricanine A can also be synthesized manually ^60^, which greatly reduces the cost of its application. Chiricanine A is plant-originating metabolic that has multiple biological functions: it can be utilized and metabolized by the recipient plant itself ^61^, which may affect its residence time within the host, an aspect that requires future research attention. As an *Aspergillus* inhibitor, chiricanine A is already widely used for eliminating *Aspergillus* spp. in the crops ^47, 62–65^, but whether *Aspergillus* spp are the unique target of chiricanine A remains unknown. We know that ST is produced by various species of *Aspergillus* ^51^, but it can also be produced by other mold species, including some members of *Penicillium*, *Emiricella*, *Chaetomium*, and *Bipolaris* genera, and most of them are pertinent to plant growth ^66, 67^. Plenty of above ST-producing microbes could be induced by PWN invasion, though in this study only *Aspergillus* was investigated. Although we show that applying chiricanine A could drastcially reduce the *Aspergillus* and ST levels in host pine trees, it is uncertain whether such chiricanine A applications can limit or benefit other microbes of host plants. In conclusion, we found a new plant-originating nematicide, chiricanine A, that has obvious advantages in terms of residence time to aid in the long-term control of PWN in pine stands, but the side effects of its and underline mechanistic action both require further attention.

## 5 CONCLUSION

Our research indicated that *Aspergillus* was able to promote PWN invasion and pathogenicity by increasing ST biosynthesis in the host plant. Further, ST accumulation triggered the biosynthesis of VOC (volatile organic compounds) that attracts JPS and drives the coexistence of PWN and JPS in the host plant, thereby encouraging the local transmission of PWD. We also found that application of an *Aspergillus* inhibitor (chiricanine A treatment) results in the absence of *Aspergillus* and decreases the in *vivo* ST amount, thereby sharply restricting the PWN development in host. Altogether, these results document, for the first time, how the function of *Aspergillus* and its metabolite ST is involved in the entire PWD transmission chain, in addition to providing a novel and long-term effective nematicide for better PWD control in the field.

## ACKNOWLEDGEMENTS

This work was supported by the Fujian Forestry Science and Technology Project (2021FKJ01, LZKG-202205) and the Forestry Peak Discipline Construction Project of Fujian Agriculture and Forestry University (72202200205). We are very grateful to the Fujian Forestry Bureau, for help in providing detailed information of the sampling areas in this study, and to Syngenta (China) Investment Co. Ltd. for technical support.

## Electronic supplementary material

The raw data of metagenomic sequencing was submitted to the NCBI bioproject database (http://www.ncbi.nlm.nih.gov/bioproject/PRJNA963075). Other related figures and tables mentioned above are included in the electronic supplementary material.

## CRediT authorship contribution statement

**JS** designed and managed the research. **JJ**, **LC**, and **WY** wrote the manuscript and **SC** performed all the analyses and drew the figures. **SS**, **XX**, **XT**, **XJ,** and **DC** carried out the field test and sampling. **YF**, **XL**, **JL,** and **YH** designed and executed the chemical treatment-related experiments.

## References

1 Zhao L, Mota M, Vieira P, Butcher RA and Sun J, Interspecific communication between pinewood nematode, its insect vector, and associated microbes. Trends in parasitology 30: 299–308 (2014).

2 Hu SJ, Ning T, Fu DY, Haack RA, Zhang Z, Chen DD, et al., Dispersal of the Japanese Pine Sawyer, *Monochamus alternatus* (Coleoptera: Cerambycidae), in Mainland China as Inferred from Molecular Data and Associations to Indices of Human Activity. PLoS One 8: e57568 (2013).

3 Hirata A, Nakamura K, Nakao K, Kominami Y, Tanaka N, Ohashi H, et al., Potential distribution of pine wilt disease under future climate change scenarios. PLoS One 12: e0182837 (2017).

4 Carnegie AJ, Venn T, Lawson S, Nagel M, Wardlaw T, Cameron N, et al., An analysis of pest risk and potential economic impact of pine wilt disease to *Pinus* plantations in Australia. Australian Forestry 81: 24–36 (2018).

5 Wu B, Liang A, Zhang H, Zhu T, Zou Z, Yang D, et al., Application of conventional UAV-based high-throughput object detection to the early diagnosis of pine wilt disease by deep learning. Forest Ecology and Management 486 (2021).

6 Li YX, Wang X, Liu ZK and Zhang XY, Research advance of pathogenic mechanism of pine wood nematode. Forest Pest and Disease 41: 11–20 (2022). (In Chinese)

7 Wang Z, Zhang Y, Wang C, Wang Y and Sung C, *Esteya vermicola* Controls the Pinewood Nematode, *Bursaphelenchus xylophilus*, in Pine Seedlings. Journal of nematology 49: 86–91 (2017).

8 Ponpandian LN, Rim SO, Shanmugam G, Jeon J, Park Y-H, Lee S-K, et al., Phylogenetic characterization of bacterial endophytes from four *Pinus* species and their nematicidal activity against the pine wood nematode. Scientific Reports 9: 12457 (2019).

9 Cai SP, Jia JY, He CY, Zeng LQ, Fang Y, Qiu GW, et al., Multi-Omics of Pine Wood Nematode Pathogenicity Associated With Culturable Associated Microbiota Through an Artificial Assembly Approach. Frontiers in Plant Science 12:798539 (2022).

10 Pires D, Vicente CSL, Menendez E, Faria JMS, Rusinque L, Camacho MJ, et al., The Fight against Plant-Parasitic Nematodes: Current Status of Bacterial and Fungal Biocontrol Agents. Pathogens 11: 1178 (2022).

11 Tian Hk, Koski TM, Zhao LL, Liu ZY and Sun JH, Invasion History of the Pinewood Nematode *Bursaphelenchus xylophilus* Influences the Abundance of *Serratia* sp. in Pupal Chambers and Tracheae of Insect-Vector *Monochamus alternatus*. Frontiers in Plant Science 13: 856841 (2022).

12 Chaudhary S, Sindhu SS, Dhanker R and Kumari A, Microbes-mediated sulphur cycling in soil: Impact on soil fertility, crop production and environmental sustainability. Microbiological Research 271: 127340 (2023).

13 Poppeliers SW, Sanchez-Gil JJ and de Jonge R, Microbes to support plant health: understanding bioinoculant success in complex conditions. Current opinion in microbiology 73: 102286–102286 (2023).

14 Feng XH, Zhang B and Sun JH, Research progress on the interaction between associated microbes and pine wood nematode-vector beetle complex. Forest Pest and Disease 41: 30–37 (2022). (In Chinese)

15 Proenca DN, Francisco R, Kublik S, Schoeler A, Vestergaard G, Schloter M, et al., The Microbiome of Endophytic, Wood Colonizing Bacteria from Pine Trees as Affected by Pine Wilt Disease. Scientific Reports 7: 4205 (2017).

16 Proenca DN, Grass G and Morais PV, Understanding pine wilt disease: roles of the pine endophytic bacteria and of the bacteria carried by the disease-causing pinewood nematode. MicrobiologyOpen 6: e00415 (2017).

17 Li Y. Response of pine endophytic bacteria flora to Bursaphelenchus xylophilus infection and Construction of insecticidal gene engineering bacteria with JK-SH007. Doctoral degree thesis, Nanjing Forestry University, Nanjing, Jiangsu, China (2018). (In Chinese)

18 Liu Y, Ponpandian LN, Kim H, Jeon J, Hwang BS, Lee SK, et al., Distribution and diversity of bacterial endophytes from four *Pinus* species and their efficacy as biocontrol agents for devastating pine wood nematodes. Scientific Reports 9: 12461 (2019).

19 Xue Q, Xiang Y, Wu XQ and Li MJ, Bacterial Communities and Virulence Associated with Pine Wood Nematode *Bursaphelenchus xylophilus* from Different *Pinus* spp. International Journal of Molecular Sciences 20: 3342 (2019).

20 Zhang HT. Preliminary study on the screening and function ofgenes related to Bursaphelenchus xylophilus by Enterobacter Ludwigii AA4. Master degree thesis, Northeast Forestry University, Harbin, Heilongjiang, China, (2021). (In Chinese)

21 Cheng XY, Xu RM and Xie BY, The role of chemical communication in the infection and spread of pine wood nematodes (*Bursaphelenchus xylophilus*). Acta Ecologica Sinica 25: 339–345 (2005). (In Chinese)

22 Teale SA, Wickham JD, Zhang F, Su J, Chen Y, Xiao W, et al., A Male-Produced Aggregation Pheromone of *Monochamus alternatus* (Coleoptera: Cerambycidae), a Major Vector of Pine Wood Nematode. Journal of Economic Entomology 104: 1592–1598 (2011).

23 Alicandri E, Paolacci AR, Osadolor S, Sorgona A, Badiani M and Ciaffi M, On the Evolution and Functional Diversity of Terpene Synthases in the *Pinus* Species: A Review. Journal of Molecular Evolution 88: 253–283 (2020).

24 Xiang Y, Wu XQ and Zhou AD, Bacterial Diversity and Community Structure in the Pine Wood Nematode *Bursaphelenchus xylophilus* and *B.mucronatus* with Different Virulence by High-Throughput Sequencing of the 16S rDNA. PLoS One 10: e0137386 (2015).

25 Fang X, Liu Y, Xiao J, Ma C and Huang Y, GC-MS and LC-MS/MS metabolomics revealed dynamic changes of volatile and non-volatile compounds during withering process of black tea. Food Chemistry 410: 135396 (2023).

26 De Vos RC, Moco S, Lommen A, Keurentjes JJ, Bino RJ and Hall RDJ, Untargeted large-scale plant metabolomics using liquid chromatography coupled to mass spectrometry. Nature protocols 2: 778–791 (2007).

27 Chen C, Gonzalez FJ and Idle JR, LC-MS-based metabolomics in drug metabolism. Drug Metabolism Reviews 39: 581–597 (2007).

28 Theodoridis G, Gika HG and Wilson ID, LC-MS-based methodology for global metabolite profiling in metabonomics/metabolomics. Trac-Trends in Analytical Chemistry 27: 251–260 (2008).

29 Bolger AM, Lohse M and Usadel B, Trimmomatic: a flexible trimmer for Illumina sequence data. Bioinformatics 30: 2114–2120 (2014).

30 Buchfink B, Xie C and Huson DH, Fast and sensitive protein alignment using DIAMOND. Nature methods 12: 59–60 (2015).

31 Li Z, Li BS, Hu ZJ, Michaud JP, Dong J, Zhang QW, et al., The ectoparasitoid *Scleroderma guani* (Hymenoptera: Bethylidae) uses innate and learned chemical cues to locate its host, larvae of the pine sawyer *Monochamus alternatus* (Coleoptera: Cerambycidae). Florida Entomologist 98: 1182–1187 (2015).

32 Hyatt D, LoCascio PF, Hauser LJ and Uberbacher EC, Gene and translation initiation site prediction in metagenomic sequences. Bioinformatics 28: 2223–2230 (2012).

33 Li W, Jaroszewski L and Godzik A, Clustering of highly homologous sequences to reduce the size of large protein databases. Bioinformatics (Oxford, England) 17: 282–283 (2001).

34 Langmead B, Trapnell C, Pop M and Salzberg SL, Ultrafast and memory-efficient alignment of short DNA sequences to the human genome. Genome Biology 10: R25 (2009).

35 Geneau CE, Waeckers FL, Luka H and Balmer O, Effects of extrafloral and floral nectar of Centaurea cyanus on the parasitoid wasp Microplitis mediator: Olfactory attractiveness and parasitization rates. Biological Control 66: 16–20 (2013).

36 Yan XF, Li XJ, Luo YQ, Xu ZC, Tian GF and Zhang TL, Taxis response of *Anoplophora glabripennis* adults to volatiles emanating from their larval host twigs. Journal of Beijing Forestry University: 80–84 (2008). (In Chinese)

37 Wang ZW, Xu HC, Zhang WW and Wang PX, *Anoplophora glabripennis* host-plant selection with main host-plant volatile chemical component analysis. Journal of Zhejiang A&F Uoiversity 33: 558–563 (2016). (In Chinese)

38 Zhang QX. Application Technical Research of Monochamus alternatus High-efficiency Attractant. Master degree thesis, South China Agricultural University, Guangzhou, Guangdong, China (2016). (In Chinese)

39 Li HM, Shen PY, Fu P, Lin MS and Moens M, Characteristics of the emergence of *Monochamus alternatus*, the vector of *Bursaphelenchus xylophilus* (Nematoda : Aphelenchoididae), from *Pinus thunbergii* logs in Nanjing, China, and of the transmission of the nematodes through feeding wounds. Nematology 9: 807–816 (2007).

40 Zhu LH, Duan HJ, Fang XY, Yang ZD, Guo CH and Wu G, Preliminary Study on Host Selection for Feeding and Oviposition of Adult Batocera horsfieldi on Walnut. Agricultural Research and Application 32: 1–4 (2019). (In Chinese)

41 Togashi K, Appleby JE, Oloumi-Sadeghi H and Malek RB, Relationship between the initial number of carried *Bursaphelenchus xylophilus* and its transmission by *Monochamus carolinensis* with reference to virulence. Nematology 24: 679–694 (2022).

42 Arias RS, Sobolev VS, Orner VA, Dang PM and Lamb MC, Potential involvement of *Aspergillus flavus* laccases in peanut invasion at low water potential. Plant Pathology 63: 354–364 (2014).

43 Ayilara MS, Adeleke BS, Akinola SA, Fayose CA, Adeyemi UT, Gbadegesin LA, et al., Biopesticides as a promising alternative to synthetic pesticides: A case for microbial pesticides, phytopesticides, and nanobiopesticides. Frontiers in Microbiology 14: 1040901 (2023).

44 Santini A and Battisti A, Complex Insect-Pathogen Interactions in Tree Pandemics. Frontiers in physiology 10: 550 (2019).

45 Guo YJ, Lin QN, Chen LY, Carballar-Lejarazu R, Zhang AS, Shao ES, et al., Characterization of bacterial communities associated with the pinewood nematode insect vector *Monochamus alternatus* Hope and the host tree *Pinus massoniana*. BMC genomics 21: 337 (2020).

46 An YB, Li YX, Ma L, Li DZ, Zhang W, Feng YQ, et al., The Changes of Microbial Communities and Key Metabolites after Early *Bursaphelenchus xylophilus* Invasion of *Pinus massoniana*. Plants-Basel 11: 2849 (2022).

47 Sobolev V, Arias R, Goodman K, Walk T, Orner V, Faustinelli P, et al., Suppression of Aflatoxin Production in *Aspergillus* Species by Selected Peanut (*Arachis hypogaea*) Stilbenoids. Journal of Agricultural and Food Chemistry 66: 118–126 (2018).

48 El-Desoky AHH, Inada N, Maeyama Y, Kato H, Hitora Y, Sebe M, et al., Taichunins E-T, Isopimarane Diterpenes and a 20-nor-Isopimarane, from *Aspergillus taichungensis* (IBT 19404): Structures and Inhibitory Effects on RANKL-Induced Formation of Multinuclear Osteoclasts. Journal of Natural Products 84: 2475–2485 (2021).

49 Cheng X, Ma FP, Yan YM, Zhao WL, Shi J, Xiao W, et al., Aspertaichunol A, an Immunomodulatory Polyketide with an Uncommon Scaffold from the Insect-Derived Endophytic *Aspergillus taichungensis* SMU01. Organic Letters 24: 7405–7409 (2022).

50 Toth L, Poor P, ordog A, Varadi G, Farkas A, Papp C, et al., The combination of *Neosartorya* (*Aspergillus*) *fischeri* antifungal proteins with rationally designed gamma-core peptide derivatives is effective for plant and crop protection. Biocontrol 67: 249–262 (2022).

51 Versilovskis A and De Saeger S, Sterigmatocystin: Occurrence in foodstuffs and analytical methods - An overview. Molecular Nutrition & Food Research 54: 136–147 (2010).

52 Hedrich R, Salvador-Recatala V and Dreyer I, Electrical Wiring and Long-Distance Plant Communication. Trends in Plant Science 21: 376–387 (2016).

53 Shen N, Wang TF, Gan Q, Liu S, Wang L and Jin B, Plant flavonoids: Classification, distribution, biosynthesis, and antioxidant activity. Food Chemistry 383: 132531 (2022).

54 Xie WF, Xu XM, Qiu WJ, Lai XL, Liu MX and Zhang FP, Expression of PmACRE1 in *Arabidopsis thaliana* enables host defence against *Bursaphelenchus xylophilus* infection. BMC Plant Biology 22: 541 (2022).

55 Takai K, Suzuki T and Kawazu K, Distribution and persistence of emamectin benzoate at efficacious concentrations in pine tissues after injection of a liquid formulation. Pest Management Science 60: 42–48 (2004).

56 Lu F, Guo K, Chen AL, Chen SN, Lin HP and Zhou X, Transcriptomic profiling of effects of emamectin benzoate on the pine wood nematode *Bursaphelenchus xylophilus*. Pest Management Science 76: 747–757 (2020).

57 Rajasekharan SK, Lee JH, Ravichandran V and Lee J, Assessments of iodoindoles and abamectin as inducers of methuosis in pinewood nematode, *Bursaphelenchus xylophilus*. Scientific Reports 7: 6803 (2017).

58 Chen Y, Zhou X, Guo K, Chen SN and Su X, Transcriptomic insights into the effects of CytCo, a novel nematotoxic protein, on the pine wood nematode *Bursaphelenchus xylophilus*. BMC genomics 22: 394 (2021).

59 Hao X, Wang BW, Chen J, Wang BY, Xu JY, Pan JL, et al., Molecular characterization and functional analysis of multidrug resistance-associated genes of Pinewood nematode (*Bursaphelenchus xylophilus*) for nematicides. Pesticide Biochemistry and Physiology 177: 104902 (2021).

60 Park BH, Lee HJ and Lee YR, Total Synthesis of Chiricanine A, Arahypin-1, trans-Arachidin-2, trans-Arachidin-3, and Arahypin-5 from Peanut Seeds. Journal of Natural Products 74: 644–649 (2011).

61 Wang HX, Qan J and Ding FY, Emerging Chitosan-Based Films for Food Packaging Applications. Journal of Agricultural and Food Chemistry 66: 395–413 (2018).

62 Sobolev VS, Localized production of phytoalexins by peanut (*Arachis hypogaea*) kernels in response to invasion by *Aspergillus* species. Journal of Agricultural and Food Chemistry 56: 1949–1954 (2008).

63 Sobolev VS, Khan SI, Tabanca N, Wedge DE, Manly SP, Cutler SJ, et al., Biological Activity of Peanut (*Arachis hypogaea*) Phytoalexins and Selected Natural and Synthetic Stilbenoids. Journal of Agricultural and Food Chemistry 59: 1673–1682 (2011).

64 Sobolev VS, Krausert NM and Gloer JB, New Monomeric Stilbenoids from Peanut (*Arachis hypogaea*) Seeds Challenged by an *Aspergillus flavus* Strain. Journal of Agricultural and Food Chemistry 64: 579–584 (2016).

65 Souto AL, Sylvestre M, Tolke ED, Tavares JF, Barbosa-Filho JM and Cebrian-Torrejon G, Plant-Derived Pesticides as an Alternative to Pest Management and Sustainable Agricultural Production: Prospects, Applications and Challenges. Molecules 26: 4835 (2021).

66 Barnes SE, Dola TP, Bennett JW and Bhatnagar D, Synthesis of sterigmatocystin on a chemically defined medium by species of *Aspergillus* and *Chaetomium*. Mycopathologia 125: 173–178 (1994).

67 Rank C, Nielsen KF, Larsen TO, Varga J, Samson RA and Frisvad JC, Distribution of sterigmatocystin in filamentous fungi. Fungal Biology 115: 406–420 (2011).

