## Supplemental Table for "The novel nematicide chiricanine A suppresses *Bursaphelenchus xylophilus* pathogenicity in *Pinus massoniana* by inhibiting *Aspergillus* and its secondary metabolite, sterigmatocystin": Data Availability Statement.docx

The raw data of metagenomic sequencing was submitted to the NCBI bioproject database (http://www.ncbi.nlm.nih.gov/bioproject/PRJNA963075). Other related figures and tables mentioned above are included in the electronic supplementary material.

1. Alicandri, E., Paolacci, A.R., Osadolor, S., Sorgona, A., Badiani, M., and Ciaffi, M. (2020). On the Evolution and Functional Diversity of Terpene Synthases in the *Pinus* Species: A Review. *Journal of Molecular Evolution* 88, 253-283. DOI: 10.1007/s00239-020-09930-8.
2. An, Y.B., Li, Y.X., Ma, L., Li, D.Z., Zhang, W., Feng, Y.Q., Liu, Z.K., Wang, X., Wen, X.J., and Zhang, X.Y. (2022). The Changes of Microbial Communities and Key Metabolites after Early *Bursaphelenchus xylophilus* Invasion of *Pinus massoniana*. *Plants-Basel* 11. 2. DOI: 10.3390/plants11212849. DOI: 10.3390/plants11212849.
3. Arias, R.S., Sobolev, V.S., Orner, V.A., Dang, P.M., and Lamb, M.C. (2014). Potential involvement of *Aspergillus flavus* laccases in peanut invasion at low water potential. *Plant Pathology* 63, 354-364. DOI: 10.1111/ppa.12088.
4. Ayilara, M.S., Adeleke, B.S., Akinola, S.A., Fayose, C.A., Adeyemi, U.T., Gbadegesin, L.A., Omole, R.K., Johnson, R.M., Uthman, Q.O., and Babalola, O.O. (2023). Biopesticides as a promising alternative to synthetic pesticides: A case for microbial pesticides, phytopesticides, and nanobiopesticides. *Frontiers in Microbiology* 14. DOI: 10.3389/fmicb.2023.1040901.
5. Barnes, S.E., Dola, T.P., Bennett, J.W., and Bhatnagar, D. (1994). Synthesis of sterigmatocystin on a chemically defined medium by species of *Aspergillus* and Chaetomium. *Mycopathologia* 125, 173-178. DOI: 10.1007/BF01146523.
6. Bolger, A.M., Lohse, M., and Usadel, B. (2014). Trimmomatic: a flexible trimmer for Illumina sequence data. *Bioinformatics* 30, 2114-2120. DOI: 10.1093/bioinformatics/btu170.
7. Buchfink, B., Xie, C., and Huson, D.H. (2015). Fast and sensitive protein alignment using DIAMOND. *Nature Methods* 12, 59-60. DOI: 10.1038/nmeth.3176.
8. Cai, S.P., Jia, J.Y., He, C.Y., Zeng, L.Q., Fang, Y., Qiu, G.W., Lan, X., Su, J., and He, X.Y. (2022). Multi-Omics of Pine Wood Nematode Pathogenicity Associated With Culturable Associated Microbiota Through an Artificial Assembly Approach. *Frontiers in Plant Science* 12. DOI: 10.3389/fpls.2021.798539.
9. Carnegie, A.J., Venn, T., Lawson, S., Nagel, M., Wardlaw, T., Cameron, N., and Last, I. (2018). An analysis of pest risk and potential economic impact of pine wilt disease to *Pinus* plantations in Australia. *Australian Forestry* 81, 24-36. https://doi.org/10.1080/00049158.2018.1440467.
10. Chaudhary, S., Sindhu, S.S., Dhanker, R., and Kumari, A. (2023). Microbes-mediated sulphur cycling in soil: Impact on soil fertility, crop production and environmental sustainability. *Microbiological Research* 271. DOI: 10.1016/j.micres.2023.127340.
11. Chen, C., Gonzalez, F.J., and Idle, J.R. (2007). LC-MS-based metabolomics in drug metabolism. *Drug Metabolism Reviews* 39, 581-597. DOI: 10.1080/03602530701497804.
12. Chen, Y., Zhou, X., Guo, K., Chen, S.N., and Su, X. (2021). Transcriptomic insights into the effects of CytCo, a novel nematotoxic protein, on the pine wood nematode *Bursaphelenchus xylophilus*. *Bmc Genomics* 22. DOI: 10.1186/s12864-021-07714-y.
13. Cheng, X., Ma, F.P., Yan, Y.M., Zhao, W.L., Shi, J., Xiao, W., Bi, E.G., and Luo, Q. (2022). Aspertaichunol A, an Immunomodulatory Polyketide with an Uncommon Scaffold from the Insect-Derived Endophytic *Aspergillus* taichungensis SMU01. *Organic Letters* 24, 7405-7409. DOI: 10.1021/acs.orglett.2c02978.
14. Cheng, X.Y., Xu, R.M., and Xie, B.Y. (2005). The role of chemical communication in the infection and spread of pine wood nematodes (*Bursaphelenchus xylophilus*). *Acta Ecologica Sinica* 25, 339-345.
15. De Vos, R.C., Moco, S., Lommen, A., Keurentjes, J.J., Bino, R.J., and Hall, R.D.J. (2007). Untargeted large-scale plant metabolomics using liquid chromatography coupled to mass spectrometry. *Nature protocols* 2, 778-791. https://doi.org/10.1038/nprot.2007.95.
16. El-Desoky, A.H.H., Inada, N., Maeyama, Y., Kato, H., Hitora, Y., Sebe, M., Nagaki, M., Kai, A., Eguchi, K., Inazumi, T., Sugimoto, Y., Frisvad, J.C., Williams, R.M., and Tsukamoto, S. (2021). Taichunins E-T, Isopimarane Diterpenes and a 20-nor-Isopimarane, from *Aspergillus* *taichungensis* (IBT 19404): Structures and Inhibitory Effects on RANKL-Induced Formation of Multinuclear Osteoclasts. *Journal of Natural Products* 84, 2475-2485. DOI：10.1021/ACS.JNATPROD.1C00486.
17. Fang, X., Liu, Y., Xiao, J., Ma, C., and Huang, Y. (2023). GC-MS and LC-MS/MS metabolomics revealed dynamic changes of volatile and non-volatile compounds during withering process of black tea. *Food Chemistry* 410. DOI：10.1016/J.FOODCHEM.2023.135396.
18. Feng, X.H., Zhang, B., and Sun, J.H. (2022). Research progress on the interaction between associated microbes and pine wood nematode-vector beetle complex. *Forest Pest and Disease* 41, 30-37.
19. Geneau, C.E., Waeckers, F.L., Luka, H., and Balmer, O. (2013). Effects of extrafloral and floral nectar of Centaurea cyanus on the parasitoid wasp Microplitis mediator: Olfactory attractiveness and parasitization rates. *Biological Control* 66, 16-20. DOI：10.1016/j.biocontrol.2013.02.007.
20. Guo, Y.J., Lin, Q.N., Chen, L.Y., Carballar-Lejarazu, R., Zhang, A.S., Shao, E.S., Liang, G.H., Hu, X., Wang, R., Xu, L., Zhang, F.P., and Wu, S.Q. (2020). Characterization of bacterial communities associated with the pinewood nematode insect vector *Monochamus alternatus* Hope and the host tree *Pinus* massoniana. *Bmc Genomics* 21. .DOI: 10.1186/s12864-020-6718-6.
21. Hao, X., Wang, B.W., Chen, J., Wang, B.Y., Xu, J.Y., Pan, J.L., and Ma, L. (2021). Molecular characterization and functional analysis of multidrug resistance-associated genes of Pinewood nematode (*Bursaphelenchus xylophilus*) for nematicides. *Pesticide Biochemistry and Physiology* 177. DOI: 10.1016/j.pestbp.2021.104902.
22. Hedrich, R., Salvador-Recatala, V., and Dreyer, I. (2016). Electrical Wiring and Long-Distance Plant Communication. *Trends in Plant Science* 21, 376-387. DOI: 10.1016/j.tplants. 2016. 01.016.
23. Hirata, A., Nakamura, K., Nakao, K., Kominami, Y., Tanaka, N., Ohashi, H., Takano, K.T., Takeuchi, W., and Matsui, T. (2017). Potential distribution of pine wilt disease under future climate change scenarios. *Plos One* 12. DOI: 10.1371/journal.pone.0182837.
24. Hu, S.J., Ning, T., Fu, D.Y., Haack, R.A., Zhang, Z., Chen, D.D., Ma, X.Y., and Ye, H. (2013). Dispersal of the Japanese Pine Sawyer, *Monochamus alternatus* (Coleoptera: Cerambycidae), in Mainland China as Inferred from Molecular Data and Associations to Indices of Human Activity. *Plos One* 8. DOI: 10.1371/journal.pone.0057568.
25. Hyatt, D., Locascio, P.F., Hauser, L.J., and Uberbacher, E.C. (2012). Gene and translation initiation site prediction in metagenomic sequences. *Bioinformatics* 28, 2223-2230. DOI: 10.1093/bioinformatics/bts429. DOI: 10.1093/bioinformatics/bts429.
26. Langmead, B., Trapnell, C., Pop, M., and Salzberg, S.L. (2009). Ultrafast and memory-efficient alignment of short DNA sequences to the human genome. *Genome Biology* 10. DOI: 10.1186/gb-2009-10-3-r25.
27. Li, H.M., Shen, P.Y., Fu, P., Lin, M.S., and Moens, M. (2007). Characteristics of the emergence of *Monochamus alternatus*, the vector of *Bursaphelenchus xylophilus* (Nematoda : Aphelenchoididae), from *Pinus* thunbergii logs in Nanjing, China, and of the transmission of the nematodes through feeding wounds. *Nematology* 9, 807-816. DOI：10.1163/156854107782331234.
28. Li, W., Jaroszewski, L., and Godzik, A. (2001). Clustering of highly homologous sequences to reduce the size of large protein databases. *Bioinformatics (Oxford, England)* 17, 282-283. DOI: 10.1093/bioinformatics/17.3.282. DOI: 10.1093/bioinformatics/17.3.282.
29. Li, Y. (2018). Response of pine endophytic bacteria *flora* to *Bursaphelenchus xylophilus* infection and Construction of insecticidal gene engineering bacteria with JK-SH007*.* Doctoral degree thesis, Nanjing Forestry University, Nanjing, Jiangsu, China.
30. Li, Y.X., Wang, X., Liu, Z.K., and Zhang, X.Y. (2022). Research advance of pathogenic mechanism of pine wood nematode. *Forest Pest and Disease* 41, 11-20. DOI：	10.19688/j.cnki.issn1671-0886.20220015.
31. Li, Z., Li, B.S., Hu, Z.J., Michaud, J.P., Dong, J., Zhang, Q.W., and Liu, X.X. (2015). The ectoparasitoid *Scleroderma guani* (Hymenoptera: Bethylidae) uses innate and learned chemical cues to locate its host, larvae of the pine sawyer *Monochamus alternatus* (Coleoptera: Cerambycidae). *Florida Entomologist* 98, 1182-1187. DOI：10.1653/024.098.0425.
32. Liu, Y., Ponpandian, L.N., Kim, H., Jeon, J., Hwang, B.S., Lee, S.K., Park, S.-C., and Bae, H. (2019). Distribution and diversity of bacterial endophytes from four *Pinus* species and their efficacy as biocontrol agents for devastating pine wood nematodes. *Scientific Reports* 9. DOI: 10.1038/s41598-019-48739-4
33. Lu, F., Guo, K., Chen, A.L., Chen, S.N., Lin, H.P., and Zhou, X. (2020). Transcriptomic profiling of effects of emamectin benzoate on the pine wood nematode *Bursaphelenchus xylophilus*. *Pest Management Science* 76, 747-757. DOI: 10.1002/ps.5575.
34. Park, B.H., Lee, H.J., and Lee, Y.R. (2011). Total Synthesis of Chiricanine A, Arahypin-1, trans-Arachidin-2, trans-Arachidin-3, and Arahypin-5 from Peanut Seeds. *Journal of Natural Products* 74, 644-649. DOI: 10.1021/np100696f.
35. Pires, D., Vicente, C.S.L., Menendez, E., Faria, J.M.S., Rusinque, L., Camacho, M.J., and Inacio, M.L. (2022). The Fight against Plant-Parasitic Nematodes: Current Status of Bacterial and Fungal Biocontrol Agents. *Pathogens* 11. DOI:10.3390/pathogens11101178.
36. Ponpandian, L.N., Rim, S.O., Shanmugam, G., Jeon, J., Park, Y.-H., Lee, S.-K., and Bae, H. (2019). Phylogenetic characterization of bacterial endophytes from four *Pinus* species and their nematicidal activity against the pine wood nematode. *Scientific Reports* 9. DOI: 10.1038/s41598-019-48745-6.
37. Poppeliers, S.W., Sanchez-Gil, J.J., and De Jonge, R. (2023). Microbes to support plant health: understanding bioinoculant success in complex conditions. *Current opinion in microbiology* 73, 102286-102286. DOI: 10.1016/j.mib.2023.102286.
38. Proenca, D.N., Francisco, R., Kublik, S., Schoeler, A., Vestergaard, G., Schloter, M., and Morais, P.V. (2017a). The Microbiome of Endophytic, Wood Colonizing Bacteria from Pine Trees as Affected by Pine Wilt Disease. *Scientific Reports* 7. DOI: 10.1038/s41598-017-04141-6.
39. Proenca, D.N., Grass, G., and Morais, P.V. (2017b). Understanding pine wilt disease: roles of the pine endophytic bacteria and of the bacteria carried by the disease-causing pinewood nematode. *Microbiologyopen* 6. DOI: 10.1002/mbo3.415.
40. Rajasekharan, S.K., Lee, J.H., Ravichandran, V., and Lee, J. (2017). Assessments of iodoindoles and abamectin as inducers of methuosis in pinewood nematode, *Bursaphelenchus xylophilus*. *Scientific Reports* 7. DOI: 10.1038/s41598-017-07074-2.
41. Rank, C., Nielsen, K.F., Larsen, T.O., Varga, J., Samson, R.A., and Frisvad, J.C. (2011). Distribution of sterigmatocystin in filamentous fungi. *Fungal Biology* 115, 406-420. DOI: 10.1016/j.funbio.2011.02.013.
42. Santini, A., and Battisti, A. (2019). Complex Insect-Pathogen Interactions in Tree Pandemics. *Frontiers in Physiology* 10. DOI: 10.3389/fphys.2019.00550.
43. Shen, N., Wang, T.F., Gan, Q., Liu, S., Wang, L., and Jin, B. (2022). Plant flavonoids: Classification, distribution, biosynthesis, and antioxidant activity. *Food Chemistry* 383. DOI: 10.1016/j.foodchem.2022.132531.
44. Sobolev, V., Arias, R., Goodman, K., Walk, T., Orner, V., Faustinelli, P., and Massa, A. (2018). Suppression of Aflatoxin Production in *Aspergillus* Species by Selected Peanut (Arachis hypogaea) Stilbenoids. *Journal of Agricultural and Food Chemistry* 66, 118-126. DOI: 10.1016/j.foodchem.2022.132531.
45. Sobolev, V.S. (2008). Localized production of phytoalexins by peanut (Arachis hypogaea) kernels in response to invasion by *Aspergillus* species. *Journal of Agricultural and Food Chemistry* 56, 1949-1954. DOI: 10.1021/jf703595w.
46. Sobolev, V.S., Khan, S.I., Tabanca, N., Wedge, D.E., Manly, S.P., Cutler, S.J., Coy, M.R., Becnel, J.J., Neff, S.A., and Gloer, J.B. (2011). Biological Activity of Peanut (Arachis hypogaea) Phytoalexins and Selected Natural and Synthetic Stilbenoids. *Journal of Agricultural and Food Chemistry* 59, 1673-1682. DOI: 10.1021/jf104742n.
47. Sobolev, V.S., Krausert, N.M., and Gloer, J.B. (2016). New Monomeric Stilbenoids from Peanut (Arachis hypogaea) Seeds Challenged by an *Aspergillus* flavus Strain. *Journal of Agricultural and Food Chemistry* 64, 579-584. DOI: 10.1021/acs.jafc.5b04753.
48. Souto, A.L., Sylvestre, M., Tolke, E.D., Tavares, J.F., Barbosa-Filho, J.M., and Cebrian-Torrejon, G. (2021). Plant-Derived Pesticides as an Alternative to Pest Management and Sustainable Agricultural Production: Prospects, Applications and Challenges. *Molecules* 26. DOI: 10.3390/molecules26164835.
49. Takai, K., Suzuki, T., and Kawazu, K. (2004). Distribution and persistence of emamectin benzoate at efficacious concentrations in pine tissues after injection of a liquid formulation. *Pest Management Science* 60, 42-48. DOI: 10.1002/ps.777.
50. Teale, S.A., Wickham, J.D., Zhang, F., Su, J., Chen, Y., Xiao, W., Hanks, L.M., and Millar, J.G. (2011). A Male-Produced Aggregation Pheromone of *Monochamus alternatus* (Coleoptera: Cerambycidae), a Major Vector of Pine Wood Nematode. *Journal of Economic Entomology* 104, 1592-1598. DOI: 10.1603/ec11076.
51. Theodoridis, G., Gika, H.G., and Wilson, I.D. (2008). LC-MS-based methodology for global metabolite profiling in metabonomics/metabolomics. *Trac-Trends in Analytical Chemistry* 27, 251-260. DOI: 10.4155/bio.12.212.
52. Tian, H.K., Koski, T.M., Zhao, L.L., Liu, Z.Y., and Sun, J.H. (2022). Invasion History of the Pinewood Nematode *Bursaphelenchus xylophilus* Influences the Abundance of Serratia sp. in Pupal Chambers and Tracheae of Insect-Vector *Monochamus alternatus*. *Frontiers in Plant Science* 13. DOI: 10.3389/fpls.2022.856841.
53. Togashi, K., Appleby, J.E., Oloumi-Sadeghi, H., and Malek, R.B. (2022). Relationship between the initial number of carried *Bursaphelenchus xylophilus* and its transmission by Monochamus carolinensis with reference to virulence. *Nematology* 24, 679-694. https://doi.org/10.1163/15685411-bja10160.
54. Toth, L., Poor, P., Ordog, A., Varadi, G., Farkas, A., Papp, C., Bende, G., Toth, G.K., Rakhely, G., Marx, F., and Galgoczy, L. (2022). The combination of *Neosartorya* (*Aspergillus*) *fischeri* antifungal proteins with rationally designed gamma-core peptide derivatives is effective for plant and crop protection. *Biocontrol* 67, 249-262. DOI: 10.1007/s10526-022-10132-y.
55. Versilovskis, A., and De Saeger, S. (2010). Sterigmatocystin: Occurrence in foodstuffs and analytical methods - An overview. *Molecular Nutrition & Food Research* 54, 136-147. DOI: 10.1002/mnfr.200900345.
56. Wang, H.X., Qan, J., and Ding, F.Y. (2018). Emerging Chitosan-Based Films for Food Packaging Applications. *Journal of Agricultural and Food Chemistry* 66, 395-413. DOI: 10.1021/acs.jafc.7b04528.
57. Wang, Z., Zhang, Y., Wang, C., Wang, Y., and Sung, C. (2017). Esteya vermicola Controls the Pinewood Nematode, *Bursaphelenchus xylophilus*, in Pine Seedlings. *Journal of Nematology* 49, 86-91. DOI: 10.21307/jofnem-2017-048.
58. Wang, Z.W., Xu, H.C., Zhang, W.W., and Wang, P.X. (2016). Anoplophora glabripennis host-plant selection with main host-plant volatile chemical component analysis. *Journal of Zhejiang A&F Uoiversity* 33, 558-563.
59. Wu, B., Liang, A., Zhang, H., Zhu, T., Zou, Z., Yang, D., Tang, W., Li, J., and Su, J. (2021). Application of conventional UAV-based high-throughput object detection to the early diagnosis of pine wilt disease by deep learning. *Forest Ecology and Management* 486. DOI：10.1016/J.FORECO.2021.118986.
60. Xiang, Y., Wu, X.Q., and Zhou, A.D. (2015). Bacterial Diversity and Community Structure in the Pine Wood Nematode *Bursaphelenchus xylophilus* and B-mucronatus with Different Virulence by High-Throughput Sequencing of the 16S rDNA. *Plos One* 10. DOI: 10.1371/journal.pone.0137386.
61. Xie, W.F., Xu, X.M., Qiu, W.J., Lai, X.L., Liu, M.X., and Zhang, F.P. (2022). Expression of PmACRE1 in Arabidopsis thaliana enables host defence against Bursaphelenchus xylophilus infection. Bmc Plant Biology 22. DOI: 10.1186/s12870-022-03929-7.
62. Xue, Q., Xiang, Y., Wu, X.Q., and Li, M.J. (2019). Bacterial Communities and Virulence Associated with Pine Wood Nematode *Bursaphelenchus xylophilus* from Different *Pinus* spp. *International Journal of Molecular Sciences* 20. DOI: 10.3390/ijms20133342.
63. Yan, X.F., Li, X.J., Luo, Y.Q., Xu, Z.C., Tian, G.F., and Zhang, T.L. (2008). Taxis response of Anoplophora glabripennis adults to volatiles emanating from their larval host twigs. *Journal of Beijing Forestry University*, 80-84.
64. Zhang, H.T. (2021). Preliminary study on the screening and function ofgenes related to *Bursaphelenchus xylophilus* by *Enterobacter Ludwigii* AA4. Master degree thesis, Northeast Forestry University, Harbin, Heilongjiang, China.
65. Zhang, Q.X. (2016). Application Technical Research of *Monochamus alternatus* High-efficiency Attractant. Master degree thesis, South China Agricultural University, Guangzhou, Guangdong, China.
66. Zhao, L., Mota, M., Vieira, P., Butcher, R.A., and Sun, J. (2014). Interspecific communication between pinewood nematode, its insect vector, and associated microbes. *Trends in Parasitology* 30, 299-308. DOI: 10.1016/j.pt.2014.04.007.
67. Zhu, L.H., Duan, H.J., Fang, X.Y., Yang, Z.D., Guo, C.H., and Wu, G. (2019). Preliminary Study on Host Selection for Feeding and Oviposition of Adult Batocera horsfieldi on Walnut. *Agricultural Research and Application* 32, 1-4.
