## Supplemental Table for "The novel nematicide chiricanine A suppresses *Bursaphelenchus xylophilus* pathogenicity in *Pinus massoniana* by inhibiting *Aspergillus* and its secondary metabolite, sterigmatocystin": supplementary figures.docx

Supplementary Material

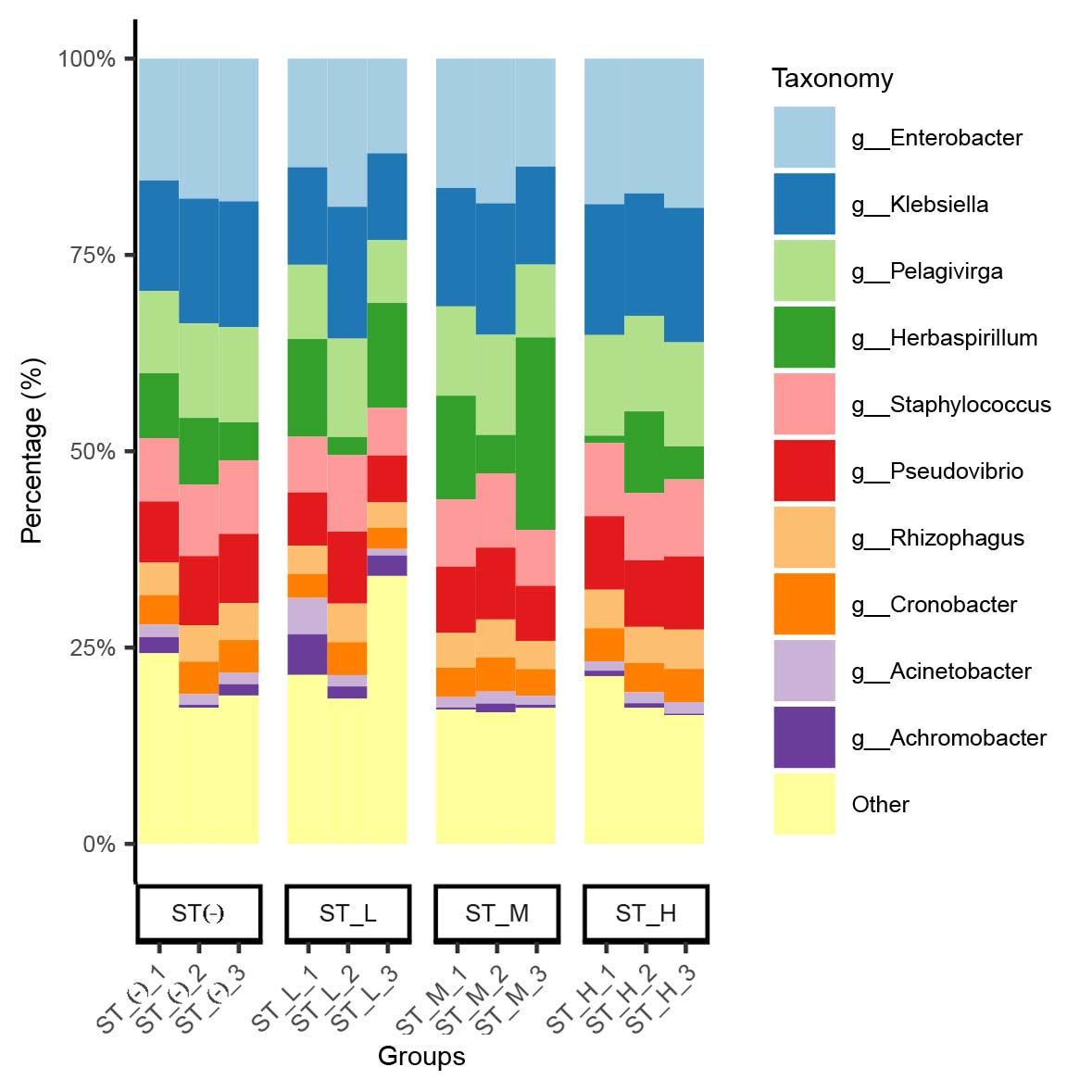

**Supplementary Figure S1. Correlations between ST and their correlated microbes in *Pinus massoniana*.**

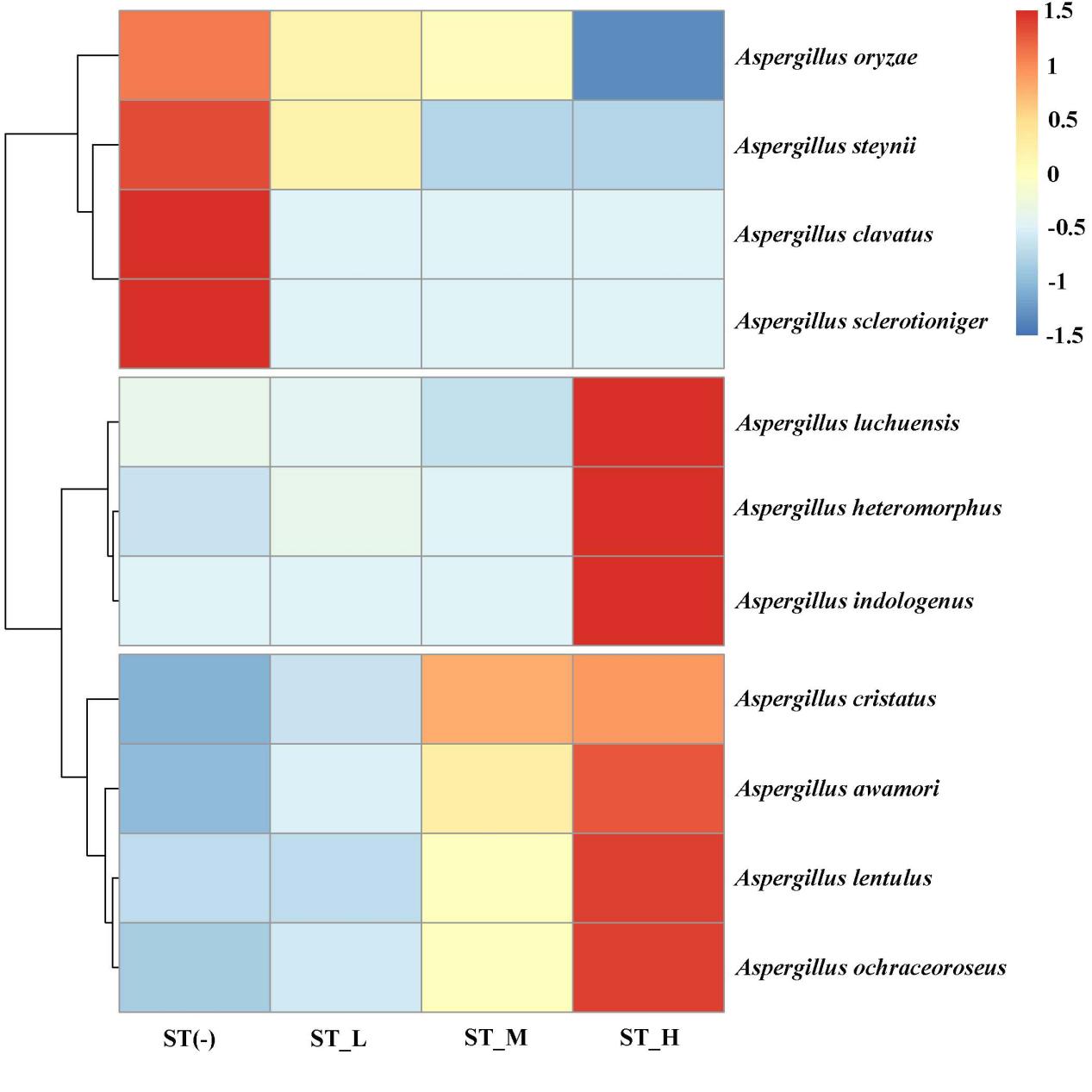

**Supplementary Figure S2. Correlations between ST and their correlated microbes of *Aspergillus* genus of fungi in *Pinus massoniana*.**
